# Lgr5+ stem/progenitor cells reside at the apex of the embryonic hepatoblast pool

**DOI:** 10.1101/485870

**Authors:** Nicole Prior, Christopher J. Hindley, Fabian Rost, Elena Meléndez Esteban, Winnie W. Y. Lau, Berthold Göttgens, Steffen Rulands, Benjamin D. Simons, Meritxell Huch

**Author notes:** Corresponding author: Meritxell Huch. These authors contributed equally (Nicole Prior, Christopher J. Hindley).

## Abstract

**Abstract:** During mouse embryogenesis, progenitors within the liver known as hepatoblasts give rise to adult hepatocyte and cholangiocyte cells. Hepatoblasts, which are specified at E8.5-E9.0, have been regarded as a homogeneous population of progenitors, which initiate differentiation into hepatocytes and cholangiocytes from E13.5 onwards. Recently, sub-populations of transcriptionally different hepatoblasts have been identified as already present at E11.5 by single cell RNAseq (scRNAseq) analysis. However, whether these transcriptional differences result from functionally heterogeneous hepatoblast populations is unknown. Here we show that the hepatoblast pool is not only transcriptionally but also functionally heterogeneous and that a sub-population of E9.5-E10.0 hepatoblasts exhibits a previously unidentified early commitment to cholangiocyte fate. Importantly, we also identify a sub-population of *bona-fide* E9.5 hepatoblasts which express the adult stem cell marker *Lgr5* and contribute to liver development by generating both hepatocyte and cholangiocyte progeny that persist for the life-span of the mouse. Using a combination of lineage tracing and scRNAseq, we show that *Lgr5* marks E9.5-E10.0 bi-potent liver progenitors residing at the apex of a hierarchy of the hepatoblast population. Notably, isolated Lgr5+ hepatoblasts can be clonally expanded *in vitro* into embryonic liver organoids, which can commit to hepatocyte or cholangiocyte fates dependent upon the culture conditions. Our study represents the first functional demonstration of heterogeneity within E9.5 hepatoblasts and identifies *Lgr5* as a marker for a sub-population of truly bi-potent liver progenitors.

**Summary Statement:** Lgr5 positive bi-potential hepatoblasts contribute to liver development and reside at the apex of an embryonic liver progenitor pool.

## Introduction

The liver is predominantly composed of hepatocytes and cholangiocytes (also known as ductal cells or biliary epithelial cells (BECs)). These epithelial cells work in conjunction with the liver stromal, endothelial and mesenchymal cells to perform essential metabolic, exocrine and endocrine functions (Zorn, 2008). In addition to performing crucial metabolic and synthetic functions, epithelial cells have a tremendous capacity for liver regeneration, which is vital to the organ due to its constant exposure to metabolic and toxic substances.

During mouse embryogenesis, liver specification from the ventral foregut endoderm begins at embryonic day (E)8.5 followed by the formation of the hepatic diverticulum. Circa E9.5, hepatic endoderm cells, the hepatoblasts, proliferate, delaminate and migrate into the adjacent septum transversum mesenchyme (STM) to form the liver bud. The hepatoblasts are the *bona-fide* embryonic progenitors for the future adult hepatocytes and cholangiocytes, whilst the STM contributes to the prospective hepatic mesenchyme (Medlock and Haar, 1983; Zorn, 2008). The STM and hepatic mesenchyme secrete several growth factors including FGF, BMP, HGF and Wnt, which promote hepatoblast proliferation, migration and survival (reviewed in Zorn, Stembook). Histological data at E13.5 shows subsets of hepatoblasts near the portal mesenchyme upregulate biliary-specific cytokeratins, indicating that biliary differentiation is initiated by E13.5 (Germain et al., 1988; Lemaigre, 2003). By contrast, hepatoblasts that are not in contact with portal veins respond to signals from the closely associated haematopoietic cells in the liver and differentiate into hepatocytes (Zorn, 2008).

Previous studies have hinted towards the bi-potential nature of hepatoblasts; initially immunohistochemical analysis in rats showed that expression of proteins such as γ-glutamyl transpeptidase, that are detected at low levels in almost all hepatoblasts, become upregulated and restricted to only differentiated cholangiocytes and not hepatocytes (Germain et al., 1988). Similarly, hepatoblasts near the portal mesenchyme destined to become cholangiocytes, transiently express *Afp* and *Alb,* two markers that later become restricted to hepatocytes (Shiojiri et al., 2001). These reports show that the hepatoblast population expresses markers of both hepatocytes and cholangiocytes, which later become lineage restricted. More recent studies have used positive selection of surface markers to isolate hepatoblasts before characterisation, as reviewed in (Miyajima et al., 2014). However, the processes that determine how hepatoblasts become fated towards cholangiocytes or hepatocytes remain largely unclear. It is also still unknown whether a single hepatoblast can give rise to both cholangiocytes and hepatocytes, *i.e.* whether single hepatoblasts are bi-potential or if there are multiple populations of unipotent hepatoblasts.

During liver development, Wnt signaling represses liver fate during endoderm patterning (McLin et al., 2007) but is required at E10 for liver bud formation (Micsenyi et al., 2004), and hepatic proliferation (Tan et al., 2008). The Wnt target gene *Lgr5* was originally described as a stem cell marker in the context of adult intestinal stem cells (Barker et al., 2007). Since then Lgr5 has been reported to be a marker of cycling adult stem cells in many other organs, such as the stomach, mammary gland and tongue, amongst others (Koo and Clevers, 2014). During development, *Lgr5* has previously been reported as a marker of bi-potent progenitors in the embryo in the context of mammary cells (Trejo et al., 2017), embryonic kidney (Barker et al., 2012) and during intestinal development (Kinzel et al., 2014). In the liver, as well as in the pancreas, *Lgr5* is not expressed under homeostatic conditions but is induced in progenitor cells in response to damage, (Huch et al., 2013a; Huch et al., 2013b). Bulk RNAseq analysis of embryonic liver tissue identified many components of the Wnt pathway, including *Lgr5,* to be differentially expressed in the developing E10.5 liver, compared to the embryonic pancreas (Rodríguez-Seguel et al., 2013). Furthermore, recent scRNAseq analysis of E11.5 livers reported that the embryonic liver harbours sub-populations of transcriptionally different hepatoblasts, some of which express *Lgr5* at the RNA level (Yang et al., 2017). However, these studies did not address whether the transcriptional heterogeneity observed at the RNA level indeed reflects a genuine functional heterogeneity of the hepatoblast pool. Also, none of these reports addressed the role of Lgr5+ cells during embryonic liver development.

Here, by combining multicolour clonal genetic lineage tracing, organoid cultures and scRNAseq analysis we demonstrate that Lgr5 marks a subpopulation of *bona-fide* bi-potential hepatoblasts, which reside at the apex of the hierarchy of an heterogenous hepatoblast pool.

## Results

### Lgr5 is a marker of hepatoblasts in the E9.5 liver

Lgr5 has been found to be expressed in the developing liver as early as E10.5 (Rodríguez-Seguel et al., 2013; Yang et al., 2017). However, these studies were performed solely at the RNA level and whether Lgr5-expressing cells are genuine hepatoblasts that contribute to liver development by generating hepatocytes and ductal cells is, at present, unknown. To formally investigate if Lgr5 marks *bona-fide* hepatoblasts, we used lineage tracing that enables the identification of all the progeny of a given cell (Kretzschmar and Watt, 2012). Hence, we generated *Lgr5-CreERT2/R26R-TdTomato* embryos, where upon tamoxifen induction, *Lgr5*+ cells and their progeny become labelled with TdTomato. Since hepatoblast delamination and formation of the liver bud occurs at E9.5 we first assessed whether Lgr5 would be expressed within this very early hepatoblast pool. To this end, we induced E9.5 embryos and collected embryos at E11.5. We found that Lgr5 expression appears as early as E9.5-E10 (considering the time for tamoxifen to induce deletion) in embryonic liver cells, as we detected TdTomato+ fluorescence in the isolated livers (Fig. 1A). We next sought to address which type of cells expressed Lgr5 during liver development. We found that at E11.5, the Lgr5+ cells labelled at E9.5 co-expressed alpha fetoprotein (AFP), a well characterised hepatoblast marker, but did not co-express markers for the endothelial or hematopoietic lineages, VEGFR3 and CD45, respectively (Fig. 1B,C). Additionally, staining with Ki67 revealed that over half of the Lgr5+ cells were proliferative (Fig. 1B,C). Collectively, these results demonstrated the existence of a population of proliferative Lgr5+ cells with hepatoblast features in the liver bud, as early as E9.5-E10.

**Figure 1:**
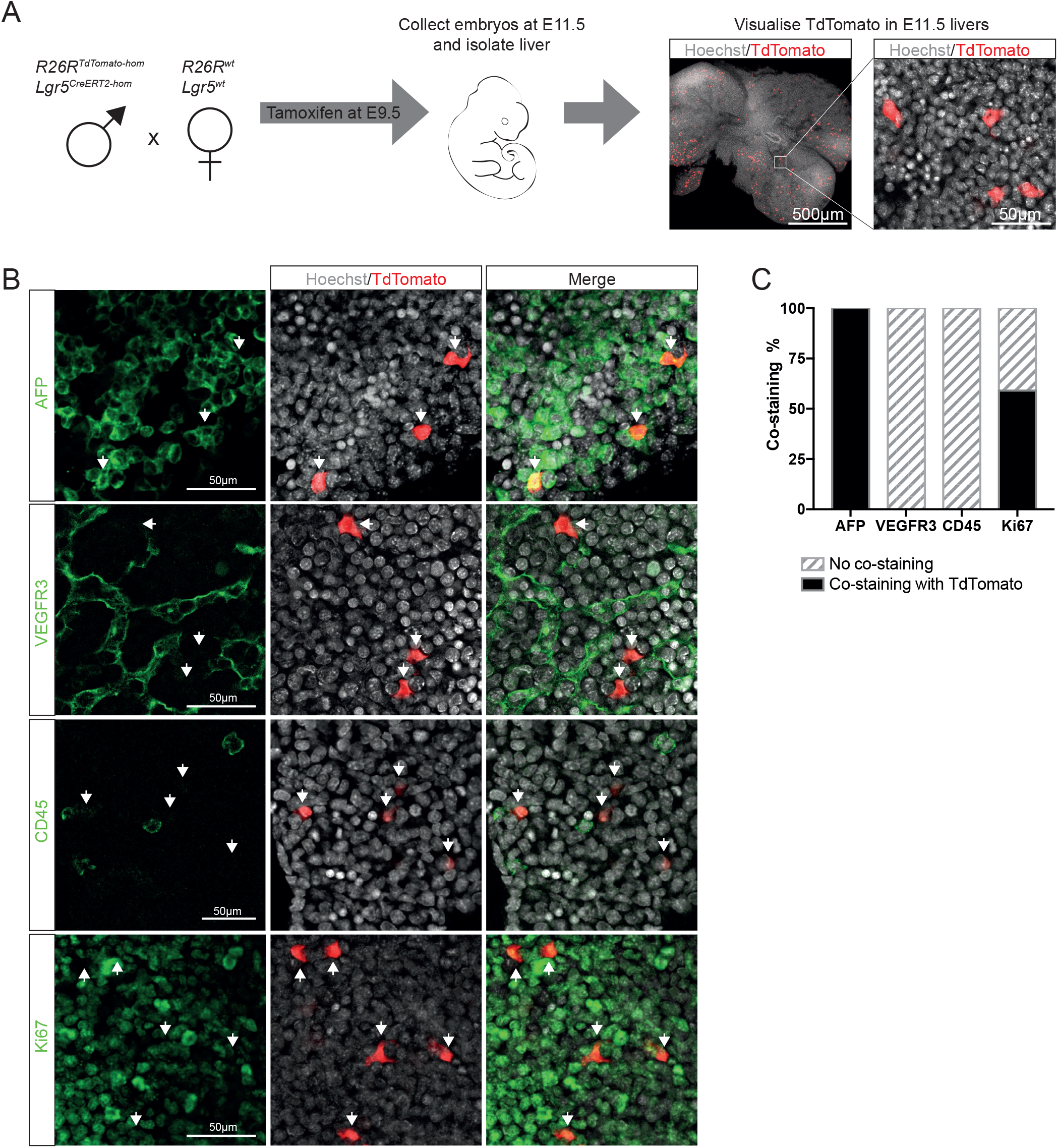
Lgr5 expression marks cells with hepatoblast features in the developing liver. (A-C) Embryos were obtained by *breeding Lgr5-ires-CreERT2^hom^; ROSA-TdTomato^hom^* males with MF1-WT females in order to generate the compound mice *Lgr5-ires-CreERT2*+/-;;*R26R-TdTomato*+/-. Administration of tamoxifen to pregnant females at E9.5 of gestation leads to activation of Cre in Lgr5+ cells and recombination at the ROSA locus to induce expression of TdTomato in Lgr5+ cells and their progeny at E9.5-E10. (A) Schematic of experimental approach. Expression of TdTomato can be detected in E11.5 livers following induction at E9.5, indicating the presence of Lgr5+ cells in the developing liver at E9.5 (n≥3 independent experiments, n=2 independent litters). Representative image of TdTomato epi-fluorescence (red) is shown. Nuclei were counterstained with Hoechst (grey). (B) Representative immunofluorescent staining of TdTomato-expressing cells co-stained with the hepatoblast marker AFP (green, top panel), endothelial marker VEGFR3 (green, middle top panel), pan-haematopoietic marker CD45 (green, middle bottom panel) and the proliferative marker Ki67 (green, bottom panel) cells. (C) Quantification of the immunostainings shown in B. Note that 100% of the Lgr5+TdTomato+ cells co-express AFP and are negative for the endothelial and haematopoietic fate markers (n>30, n=2 independent litters). Half of the TdTomato-expressing cells are proliferative (Ki67+, n>50, n=2 independent litters).

To assess if these Lgr5+ cells are *bona-fide* hepatoblasts, we analysed their contribution to the formation of both mature hepatocytes and cholangiocytes in the postnatal liver. To test this, we induced *Lgr5-CreERT2/R26R-TdTomato* embryos at E9.5 and collected the livers postnatally, over the course of a year (Fig. 2A). We detected TdTomato+ descendants of the initially labelled E9.5-E10 Lgr5+ cells at all timepoints analysed (from 1 month up to 1 year after birth) in all three functional zones of the liver (zones 1-3) (Fig. 2B). Importantly, we found both hepatocytes as well as ductal cells (cholangiocytes), as descendants of the E9.5 Lgr5+ hepatoblasts. Remarkably, induction at later time points (E13.5) resulted in only hepatocyte labelling at 1 month after birth, which indicated that, by E13.5, Lgr5+ embryonic liver cells are already hepatocyte-fated hepatoblasts (Fig. S1A,B). Of note, induction at earlier timepoints (E7.5 and E8.5) did not result in any labelled progeny in the postnatal liver (Fig. S1A), suggesting that embryonic Lgr5 exclusively marks liver progenitors after specification and liver bud formation, but not definitive endoderm or foregut progenitors that will contribute to the prospective liver tissue. No labelling was detected in non-tamoxifen induced mice (Fig. S1C). Collectively, our lineage tracing results demonstrate that Lgr5 is indeed a *bona-fide* hepatoblast marker that identifies E9.5-E10 liver bud hepatic progenitors with the capacity to give rise to adult hepatocytes and cholangiocytes.

**Figure 2:**
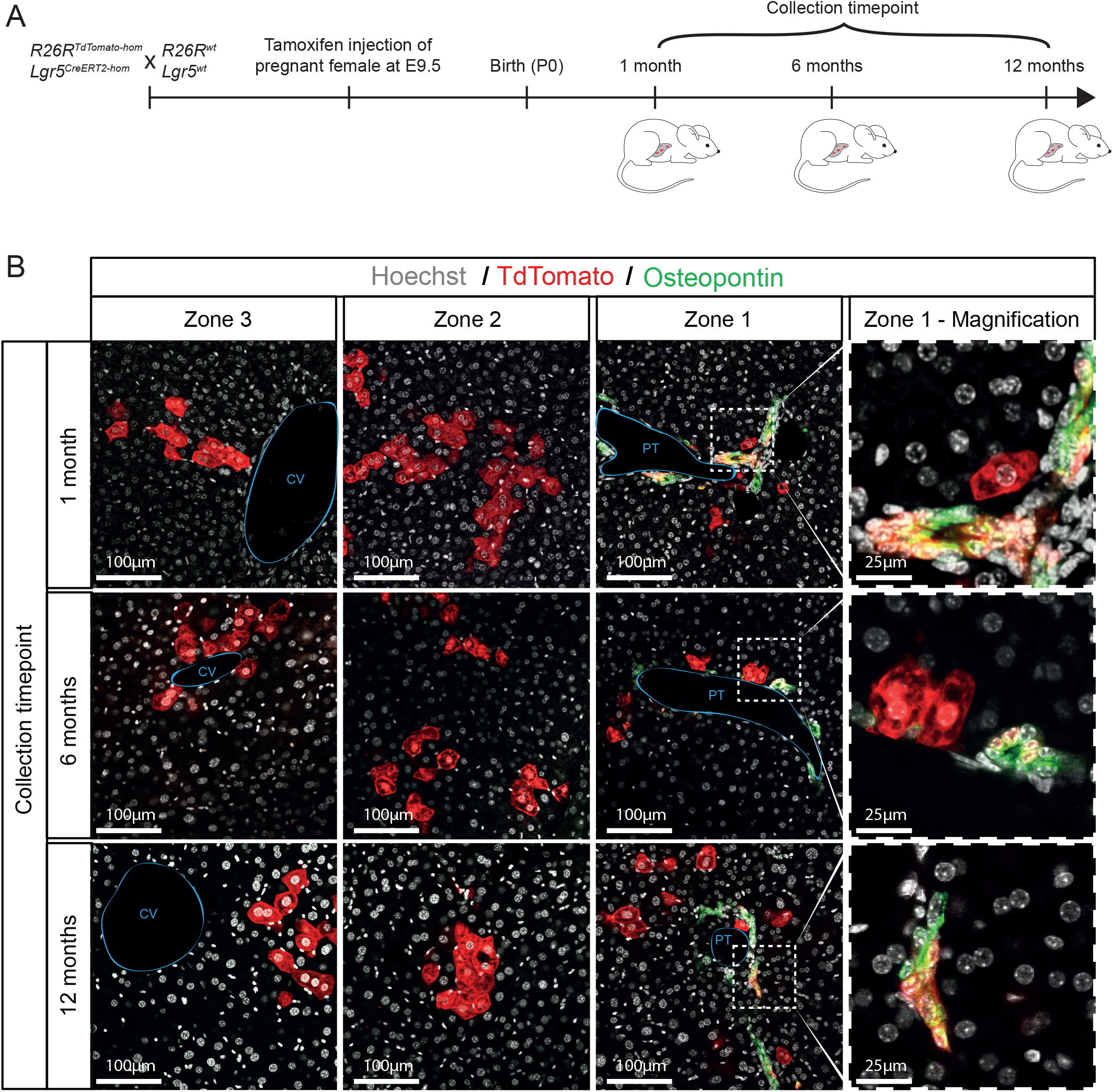
Lgr5 is a marker of *bona-fide* hepatoblasts in vivo. (A-B) *Lgr5-ires-CreERT2^hom^; ROSA-TdTomato^hom^* males were breed with MF1-WT females in order to generate the compound mice *Lgr5-ires-CreERT2*+/-;*R26R-TdTomato*+/-. Cre activity was induced in *Lgr5-ires-CreERT2*+/-;*R26R-TdTomato*+/-embryos at E9.5 by injecting pregnant females with tamoxifen (1-2mg) as described in methods. Livers were collected at the indicated postnatal timepoints and processed for immunofluorescence analysis to identify the progeny of Lgr5+ E9.5 hepatoblasts in the postnatal liver. (A) Schematic of experimental approach. (B) Lgr5+ descendants (Tdtomato+ cells, red) include both hepatocytes and ductal cells (Osteopontin, green). E9.5 Lgr5 progeny are found distributed along all 3 zones of the liver lobule; the portal triad (PT, zone 1), the central vein (CV, zone 3) and in the intermediate region (zone 2) at all time points analysed, up to 12 months after birth. In the portal area labelled cells include both hepatocytes and ductal cells, indicating that the E9.5-E10 induced Lgr5+cells were *bona-fide* hepatoblasts. Right panels represent a magnified area from Zone1.

### Embryonic Lgr5+ cells are *bona-fide* bi-potential hepatoblasts *in vivo*

To date it has not been clear whether hepatoblasts are truly bipotential *i.e.* whether a single hepatoblast gives rise to both cholangiocytes and hepatocytes, or conversely whether hepatoblasts are unipotent and two types of hepatoblasts, progenitors for either hepatocyte or cholangiocyte fates, coexist within the hepatoblast population. To formally assess whether our E9.5-E10 Lgr5+ hepatoblasts are indeed bi-potential, we opted to use a multicolour lineage tracing approach, where single cell clones are labelled with different colours in order to discriminate the contribution of each individual clone to a given fate. Therefore, we used lineage tracing with the R26R-Confetti reporter in combination with Lgr5-CreERT2, where following clonal induction with tamoxifen at E9.5, Lgr5+ cells are labelled stochastically with either RFP, YFP, GFP or CFP (Snippert et al., 2010)(Fig. 3A). As expected, we detected distinct clones labelled with one of the four fluorescent proteins at all time points analysed (P0-P17) (Fig. 3B).

**Figure 3:**
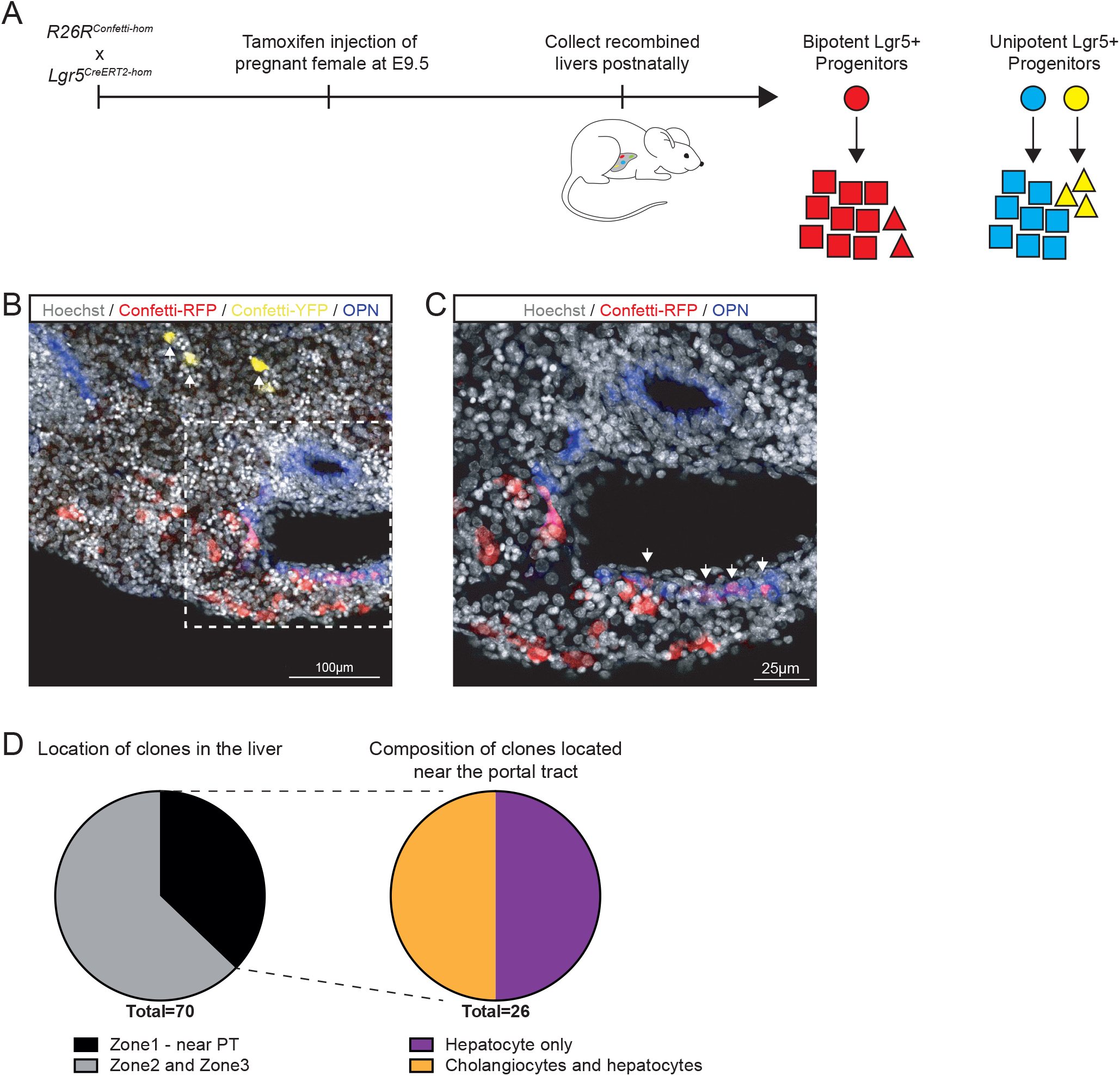
E9.5 Lgr5+ hepatoblasts are bi-potential. (A-D) To assess whether a single Lgr5+ hepatoblast is bi-potent, i.e., is able to give rise to both hepatocytes and ductal cells we generated the compound mouse *Lgr5-ires-CreERT2;R26R-Confetti* by breeding the *Lgr5-ires-CreERT2^hom^* with the multicolour Confetti reporter R26R-Confettihom which results in cells labelled in one of four colours (RFP, YFP, mCFP, nGFP) when induced with 0.16mg/g tamoxifen at E9.5 (see Supplementary Fig. 2 and Supplementary Dataset 2 for details). In this way we can identify individual clones to assess the bi-potentiality of individual hepatoblasts. (A) Schematic of experimental design. The scheme illustrates the two potential outcomes from the experiment; a single Lgr5+ hepatoblast (red circle) is bi-potent and gives rise to both hepatocytes (red squares) and ductal cells (red triangles) or, alternatively, single Lgr5+ hepatoblasts (blue and yellow circles) are unipotent and independently give rise to hepatocytes (blue squares) or ductal cells (yellow triangles). (B-C) Representative images of a P0 *Lgr5-CreERT2+/-; R26R-Confetti+/-* liver following induction at E9.5. Ductal cells were co-stained with OPN (blue, white arrows). Nuclei were stained with Hoechst. (B) Low power magnification of a liver section showing two labelled clones, a red clone and a yellow clone (white arrows). (C) Magnification showing that the red clone contains both hepatocytes and ductal cells (white arrows). (D) Pie-charts showing the total number of clones identified (n=70) and the fraction of these that are located in the portal area (n=26). Note that from the total number of clones found on the portal area, half of them (n=13) contained both hepatocytes and ductal cells of the same colour. At the induction dose used, merging of clones of the same colour occurs with a 1.2% probability (see Supplementary Fig. 2B), which confirms that at least 11 of the 13 bi-potent clones identified arise from a single Lgr5+ cell, demonstrating that indeed Lgr5+ cells are bi-potent at E9.5. Experiments were performed in n=3 embryos.

We recently showed that clonal fragmentation due to the expansion of the tissue during development results in dispersion of clones throughout the tissue, hence imposing an additional constraint on clonal fate studies during development (Rulands et al., 2018). Therefore, to ensure that we were scoring cells within individual clones, we opted to only score as positive bi-potential clones, those clones in the portal tract where ductal cells and hepatocytes were juxtaposed. We scored 70 individual clones, 81% of which were comprised only of hepatocytes, whereas no cholangiocyte-only clones were found. Crucially, from all the clones identified, 37% were identified near a portal tract and half of these (50%) contained both labelled hepatocytes and cholangiocytes with the same confetti colour (Fig. 3C,D)(Movie S1). As, expected, we also found clones formed of hepatocytes without cholangiocytes throughout the liver in zones 2 and 3. These results are not explained by clone merging, i.e., the probability of labelled cells with the same colour counted as a single clone but originating from two recombination events, since our quantification of merging events revealed a 1% probability (p=0.012) (Fig. S2A,B), indicating that from the 13 bi-potential clones identified at least 11 clones are truly bi-potent. As before, no labelling was detected in non-tamoxifen induced mice (Fig. S2C).

Therefore, the clonal analysis of Lgr5+ hepatoblasts demonstrates that *Lgr5* marks a population of E9.5-E10 hepatoblasts in which at least some are fully bi-potential at E9.5 time point.

### Lgr5+ embryonic liver cells grow into organoids *in vitro* and generate both ductal and hepatocyte fated organoids in culture

In the adult liver of mouse (Huch et al., 2013b) and human (Huch et al., 2015), single Lgr5+ cells grown clonally into cholangiocyte liver organoids retain the bi-potential characteristics of adult liver cholangiocyte progenitors, being able to self-duplicate while retaining the capacity to differentiate into both hepatocytes and ductal cells *in vitro.* To assess whether our Lgr5+ embryonic liver bi-potent hepatoblast would also be able to retain the self-renewing and bi-potent differentiation characteristics in culture if grown into organoids, we sought to establish liver organoids from Lgr5+ hepatoblasts and assess their bi-potential characteristics.

Recently, further optimization of our protocols to expand human adult liver cells (Huch et al., 2015) have facilitated the expansion of human embryonic (week 11-20 human gestation) liver tissue as 3D organoid cultures (Hu et al., 2018). However, the medium requirements to establish mouse liver organoids from mouse embryonic bi-potent hepatoblasts have not yet been reported. Hence, we first sought to establish culture conditions that would enable the expansion of mouse organoid cultures from the embryonic liver. For that, we first opted to use the entire liver tissue isolated from E10.5 mouse embryonic livers (without selection for specific hepatoblast cells) and test several conditions in order to establish both cholangiocyte and hepatocyte organoids from these early progenitors. We opted to use E10.5 instead of E9.5 embryos for practical reasons; at E9.5 the prospective liver has not yet formed a clear organ structure and therefore it was not possible to isolate only the liver, which resulted in contamination from other foregut derived tissues, especially stomach (data not shown). To establish cholangiocyte organoids from the embryonic liver we optimized our previously published protocol to expand mouse adult liver organoids (Broutier et al., 2016; Huch et al., 2013b) by adding TGFb inhibitor and Forskolin (Fig. S3A). In parallel, to establish hepatocyte organoids, we adapted the recently published protocol from human embryonic liver (Hu et al., 2018) by removing FGF7 during passaging (Fig. S3B). Using these culture conditions we could expand mouse embryonic liver organoids for up to 5 passages (3 months in culture).

Next, we assessed whether single Lgr5+ cells isolated from E10.5 livers would retain their ability to differentiate into either lineage *in vitro* when cultured in our optimized cholangiocyte and hepatocyte media conditions. To this end, we first established a sorting strategy that would enable isolation of pure populations of Lgr5+ E10.5 hepatoblasts, utilising the Lgr5-GFP mouse line, where the eGFP reporter is knocked-in into the Lgr5 locus (Barker et al., 2007), combined with co-staining with anti-Liv2, which specifically labels liver progenitors from E9.5-E13.5 (Nierhoff et al., 2005; Watanabe et al., 2002). We confirmed the hepatoblast nature of E10.5 Lgr5+ liver progenitors by co-staining with anti-Liv2 (Fig. 4A). Given that the developing liver serves as the site of haematopoiesis from E10.5 until the perinatal stage (Sasaki and Sonoda, 2000), we used negative selection of the hematopoietic marker CD45 and endothelial marker CD31 to limit contamination by non-liver progenitor cells. Embryonic livers were collected at E10.5-E12.5, enzymatically digested and then fluorescence activated cell sorting (FACS) was used to isolate liver progenitors and also Lgr5+ hepatoblasts (Fig. S4A). The sorting strategy was sequentially gated based on cell size, singlets, Liv2^+^/CD31^−^/CD45^−^/Lgr5-GFP^+^. Sorted Lgr5^+^ cells were embedded in Matrigel (as extracellular matrix) and cultured under our two optimized medium conditions (Fig. 4B). The morphology of structures generated was dependent on the culture medium used. Addition of ‘duct’ medium, resulted in the generation of single-layer epithelial spheres (Fig. 4C). The duct-like morphology of embryonic organoids cultured with ‘duct’ medium are reminiscent of adult mouse liver organoids (Huch et al., 2013b). These cultures expressed the classic cholangiocyte marker-*Krt19* (Fig.4E). Conversely, Lgr5+ cells cultured with ‘hep’ medium resulted in more compacted, dense and budding, bunch-of-grapes shape, structures (Fig. 4D), that resembled the recently published human embryonic hepatocyte organoids (Hu et al., 2018). The hepatocyte nature of the embryonic cells grown with ‘hep’ medium was confirmed by gene expression analysis, which showed high levels of *Albumin* in the embryonic cells with very low levels of *Krt19* (Fig. 4E). Furthermore, immunofluorescence analysis of the embryonic cells cultured with ‘hep’ medium confirmed also their hepatocyte nature, with clear expression of the hepatocyte marker HNF4a and no detection of the ductal marker EpCAM (Fig. 4F). As expected, they also express *Afp,* a classic hepatoblast marker (Fig. 4G), which is absent in our adult-derived organoids (Fig. S3C). Therefore, these results confirm that Lgr5+ hepatoblasts retain self-renewal and differentiation capacity, being capable of differentiating towards both cholangiocyte and hepatocyte fates *in vitro,* when grown in our 3D organoid culture conditions.

**Figure 4:**
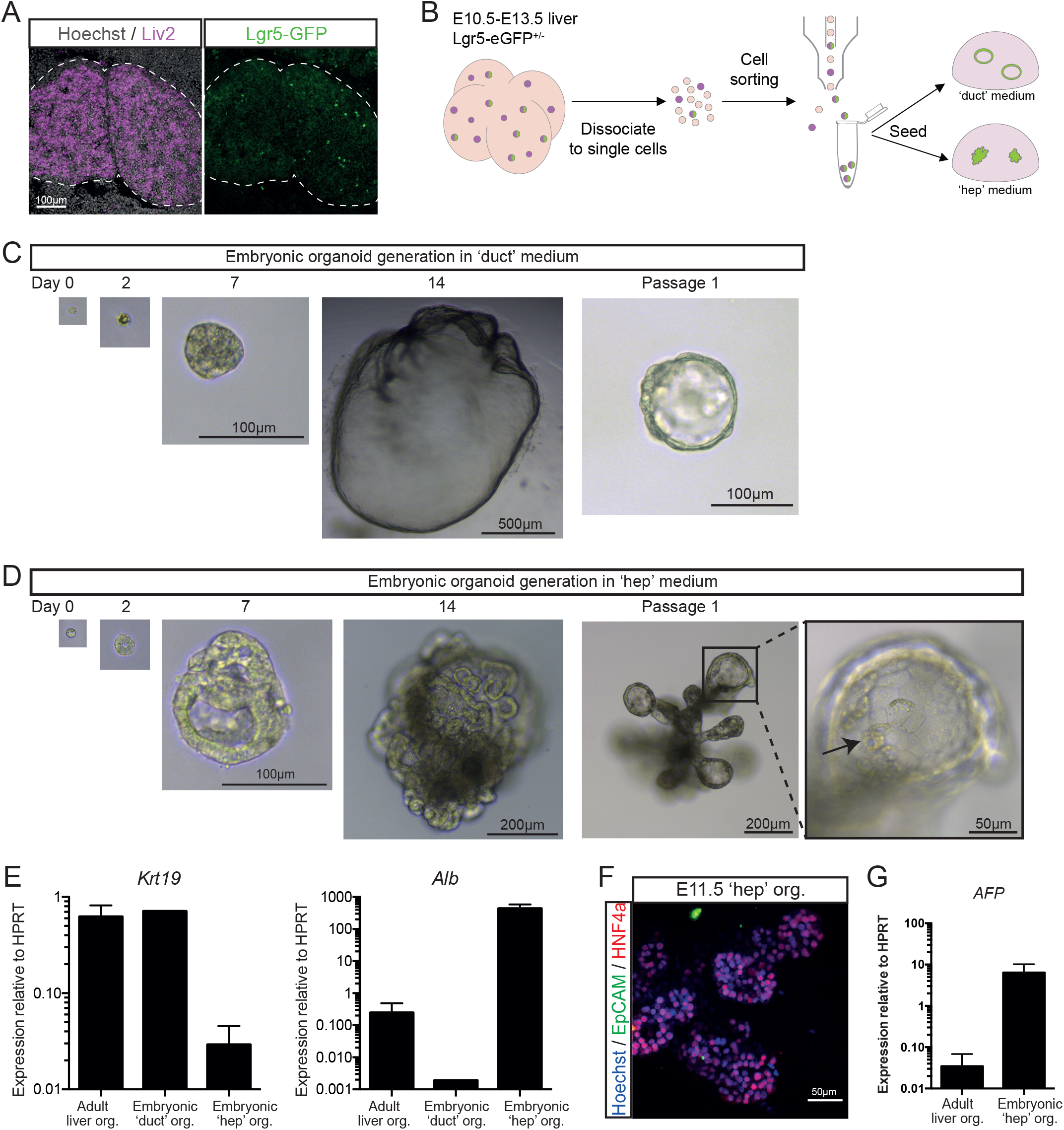
Lgr5+ embryonic liver cells grow into organoids that generate both ductal and hepatocyte fated cells in vitro. (A) Low power magnification of a section of an E10.5 liver showing co-labeling of Lgr5-eGFP+ cells (green) with the liver progenitor marker Liv2 (purple). Nuclei were counterstained with Hoechst. (B-F) Embryonic liver organoids were generated from FACS sorted Lgr5-eGFP positive hepatoblasts obtained from E10.5-E13.5 *Lgr5-EGFP-IRES-creERT2.* Sorted Liv2+/CD31-/CD45-/Lgr5-eGFP+ cells were embedded in Matrigel and cultured in our optimized duct or hepatocyte (hep) mouse embryo liver medium as described in methods and supplementary figure S3A,B. (B) Schematic of the experimental approach. (C-D) Representative image of a mouse embryo liver organoid derived from a single Lgr5-eGFP+ cell and cultured in cholangiocyte medium (C) or hepatocyte medium (D). Note the hepatocyte morphology on the organoids grown in hepatocyte medium (D). (E) Expression of the ductal marker Krt19 is mainly detected in the ductal organoids grown in ductal medium and in control adult liver organoids, while the hepatocyte marker Albumin (Alb) is only detected in embryonic cells cultured in hepatocyte medium (‘hep’). Graphs represent mean ± SEM of n≥2 experiments. (F) Immunofluorescence staining for the ductal marker EpCAM and the hepatocyte marker HNF4 in hepatocyte organoids derived from Lgr5-eGFP+ E11.5 embryonic liver cells. Note that the organoids grow as solid structures with all cells marked by HNF4a while the ductal marker EpCAM is virtually not detected. Adult liver organoids which express high levels of the ductal marker EpCAM were used as positive controls for the staining (see supplementary figure S3C). (G) embryonic cells cultured in ‘hep’ expressed the hepatoblast marker AFP whilst adult ductal organoids do not. Graphs represent mean ± SEM of n≥2 experiments.

### scRNAseq identifies distinct heterogeneity within the hepatoblast population

To address whether all hepatoblasts express Lgr5 or whether instead Lgr5 is a marker of a specific sub-population of *bona-fide* bipotential hepatoblasts, we performed single cell RNA sequencing (scRNAseq) analysis on both Lgr5+ hepatoblasts and bulk embryonic liver cells derived-from either E10.5 or E13.5 livers. To isolate liver progenitors (Liv2+) and also Lgr5+ hepatoblasts (Liv2+Lgr5+), we applied our established sorting strategy to E10.5 and E13.5 embryonic livers derived from Lgr5-GFP mice (Fig. 5A). Therefore, sorted Liv2^+^, CD31^−^ /CD45^−^ (Liv2+, bulk hepatoblast population) and Liv2^+^/CD31^−^/CD45^−^Lgr5-GFP^+^ (Lgr5^+^ hepatoblasts) were used for single cell RNA sequencing analysis based on the Smart-seq2 protocol (Picelli et al., 2014). scRNAseq analysis was conducted on 943 sorted cells. Following quality control, 653 cells were taken for further analysis. To reduce technical variability between biological replicates, we applied batch effect correction by matching mutual nearest neighbours.

**Figure 5:**
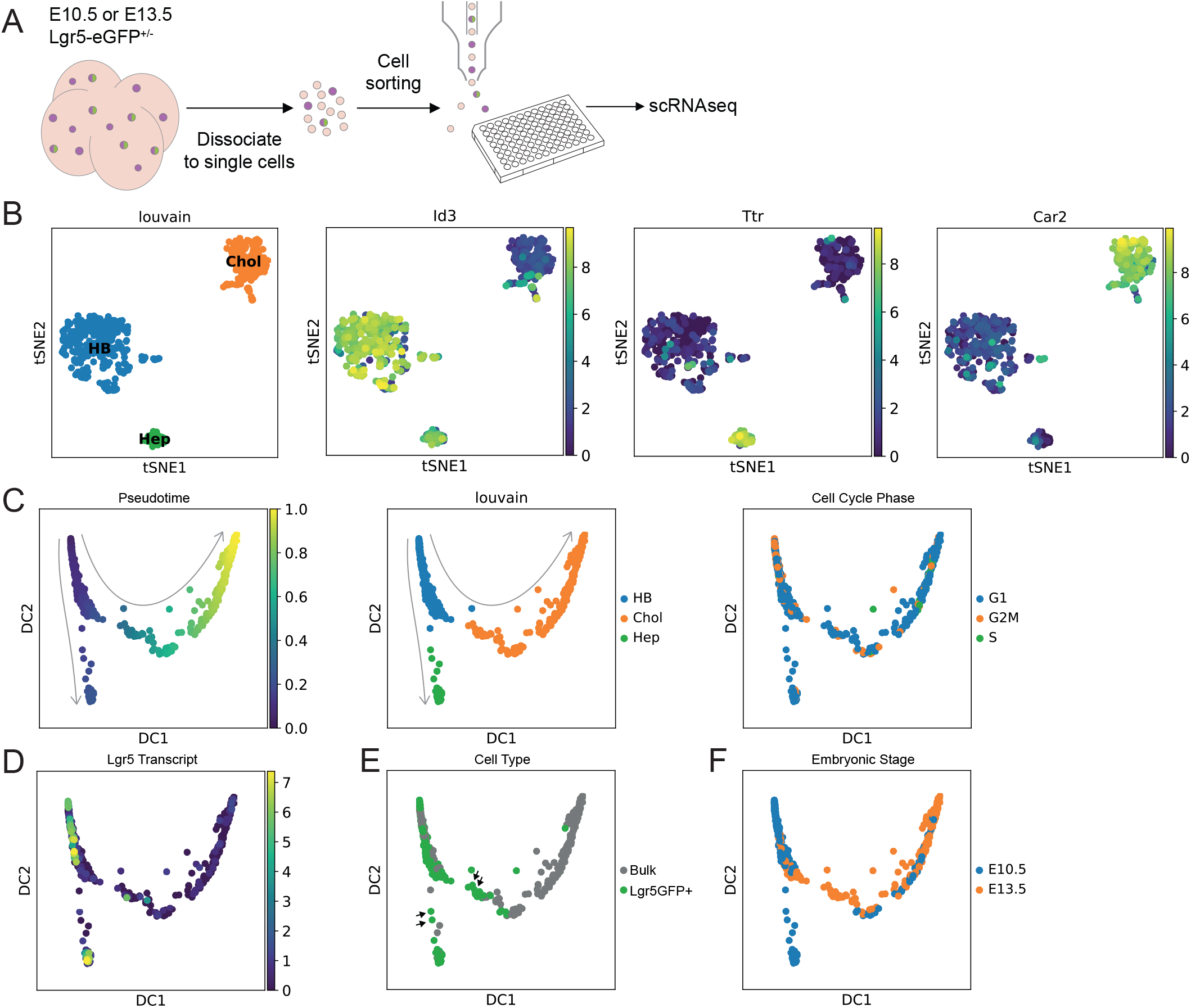
scRNAseq of hepatoblasts reveals heterogeneity in the hepatoblast population. (A-D) A bulk population of hepatoblasts (Liv2+) or Lgr5-eGFP positive hepatoblasts (Liv2+ and Lgr5-eGFP+) from E10.5-E13.5 *Lgr5-EGFP-IRES-creERT2* embryos were obtained by FACS sorting using the using the combination: *Liv2+/CD31-/CD45-/Lgr5-eGFP+* cells as described in Supplementary Fig.3. Both, hepatoblasts cells (Liv2+, bulk hepatoblast pool) and Lgr5+ hepatoblasts (Liv2+,eGFP+) were processed for scRNAseq analysis using Smartseq2 protocol. Number of clusters identified and subsequent analysis was performed as described in methods. (B) Clustering analysis (louvain clustering) of all the cells analysed (653 sorted cells from E10.5 and E13.5 embryos) classified all the cells into 3 different clusters: an hepatoblast cluster that exhibits features of hepatoblasts only and that we name “early hepatoblast cluster” (HB, blue); an hepatoblast cluster with hepatocyte-like features (‘Hep’, green) and an hepatoblast cluster with cholangiocyte-like features (‘Chol’, orange). Representative marker genes of each of these three clusters are shown: *Id3* (‘HB’ cluster), *Ttr,* hepatocyte-like cluster and *Car2,* cholangiocyte-like cluster. Clusters are represented here using tSNE plots. (C) Diffusion pseudotime analysis of E10.5 and E13.5 cells shows the early hepatoblast cluster precedes the divergence of hepatocyte-like cells or cholangiocyte-like cells and has a higher proportion of cells in G2M phase. Left panel, diffusion map showing DC1 and DC2 components; middle panel, diffusion map where the 3 clusters identified by louvain clustering are shown; right panel, diffusion map where the cell cycle phases are shown. Arrows represent developmental trajectory originating from the proliferating hepatoblast cluster. (D-F) Segregation of the data by expression of Lgr5 transcript (D) or by sorting for Lgr5-eGFP (E) or by time point (F). (D) Lgr5 transcript levels as determined using single cell sequencing superimposed on the pseudotime analysis of all the cells (both from the Liv2+ bulk as well as from both time points). (E) Lgr5-eGFP+ cells based on the FACS data superimposed on the pseudotime analysis of all cells (Liv2+, bulk, grey; Liv2+Lgr5-eGFP+, green). Note that there are some Lgr5-eGFP+ cells that were sorted as GFP+ but that have down-regulated the Lgr5-transcript (black arrows), indicating that these are immediate descendants of the Lgr5+ cells. (F) Diffusion map showing the cells segregate by time point (blue, E10.5; orange, E13.5). Note that cells sorted at E10.5 are found in the proliferating hepatoblast cluster, in the cells moving towards the hepatocyte-like cluster, the hepatocyte-like cluster and cells located in the cholangiocyte-like cluster. At E13.5 the sorted cells map to the hepatoblast cluster, the cells moving towards the cholangiocyte-like cluster and the cholangiocyte-like cluster. Sorted cells no longer map to the hepatocyte cluster, which may indicate that hepatocyte-committed hepatoblasts do not express the Liv2 epitope at E13.5.

To define embryonic liver progenitor populations, we performed dimensionality reduction using t-distributed stochastic neighbour embedding (tSNE) analysis on all 653 cells (Fig. 5B). This identified three distinct progenitor populations which were confirmed by Louvain clustering. These three clusters signified biological differences, since each cluster contained cells from each biological replicate. The biological differences were confirmed by the expression of distinct marker genes (Fig. 5B, S4A and Supplementary dataset 1). The cell type identity of each cluster was assigned based on examining the marker genes and comparing them to publicly available gene expression patterns in human or mouse liver (Broutier et al., 2017; Yang et al., 2017). We found that the three clusters corresponded to proliferating hepatoblasts (HB), hepatocyte-like progenitors (Hep) and cholangiocyte-like progenitors (Chol), which express higher levels of representative markers. The hepatoblast cluster contained *Id3, Mdk* and *Gpc3,* all described as hepatoblast markers (Su et al., 2017; Yang et al., 2017) while the hepatocyte-like cluster presented *Ttr, Alb, Apoa1, Apoa2* and *C3* all known hepatocyte markers and the cholangiocyte-like cluster expressed the ductal cell genes *Car2, Cd44* and *Bcl11a* (Yang et al., 2017) (Fig. 5B and Supplementary dataset 1_S1-S7). Of note, within the E10.5 cholangiocyte-like cluster we found two sub-clusters (Supplementary dataset 1_S7). Although we found known cholangiocyte and hepatocyte markers, new markers for these clusters were also revealed by the analysis (Fig. S5A and Supplementary dataset 1_S1-S7).

To establish developmental trajectories between the different cells of the 3 clusters we calculated diffusion maps and diffusion pseudotime. This analysis revealed a developmental trajectory originating from the proliferating hepatoblast cluster which bifurcated towards either the hepatocyte-like cluster or cholangiocyte-like cluster (Fig. 5C). We found the proliferating hepatoblast cluster contained a higher proportion of cells in G2M phase, indicating an increased number of proliferative cells (Figs. 5C,S5B). When analysing the lineage trajectories we took advantage of the Lgr5-GFP mouse line (Barker et al., 2007), which enabled us to identify cells that expressed Lgr5 RNA via sequencing and compare whether they were GFP positive from the FACS (Fig. 5D,E). Since the GFP protein is more stable than the transcript, we used the comparison between the Lgr5GFP sorted cells and the cells expressing Lgr5 transcript as a proxy to identify the immediate descendants of Lgr5+ cells in the scRNAseq population. Notably, most of the Lgr5+cells mapped to the hepatoblast cluster, representing 2% of the total number of Liv2+ hepatoblasts at E10.5 (Fig. S5C). Interestingly, we observed that as cells exit the hepatoblast cluster and become committed to either of the two epithelial lineages Lgr5 transcript levels decrease (Fig. 5D). Many of these transitioning cells (Fig. 5E, black arrows) were negative for Lgr5 transcript but positive for Lgr5GFP, indicating that these cells have only recently reduced Lgr5 levels as the GFP protein has not yet degraded and can be considered immediate descendants of the Lgr5+ pool. Interestingly, once cells have transitioned to the hepatocyte cluster *Lgr5* is upregulated whilst *Lgr5* expression is not reinitiated in the cholangiocyte cluster.

Segregation of the data by embryonic stage shows that E10.5 cells contribute to the proliferating hepatoblast cluster, the intermediate cells that are moving from the hepatoblast towards the hepatocyte-like cluster, the hepatocyte-like cluster and cells located at the far end of the cholangiocyte-like cluster (Fig. 5F). At E10.5, though, we find very few cells in the transition between the proliferating hepatoblast and cholangiocyte-like cluster, however, some of them were Lgr5GFP+ that had downregulated Lgr5 transcript, suggesting that they were immediate descendants of the E10.5 Lgr5+ proliferating hepatoblast cluster cells. Similarly, cells occupying this intermediate space were readily identified at E13.5, the majority of which also appear to have recently downregulated *Lgr5,* again indicating that they were immediate descendants of the Lgr5+ cells on the hepatoblast cluster. This implies that the proliferating Lgr5+ hepatoblasts do indeed give rise to cholangiocytes at E10.5 but with higher proportion at E13.5. Intriguingly, E13.5 cells (either Lgr5GFP+ or bulk) significantly contributed to the cholangiocyte-like cluster, but we did not find E13.5 cells that mapped to the hepatocyte cluster (Fig. 5F). This result was in striking disagreement with our knowledge of liver development and our E13.5 lineage tracing results from *the Lgr5-CreERT2* allele, which provides evidence that E13.5 Lgr5+ tracing resulted in labelling of only hepatocytes (Fig. S1B), indicating that at E13.5, cells committed to a hepatocyte fate are indeed present and express *Lgr5.* Our interpretation of this discrepancy between the lineage tracing and the scRNAseq data is that the cells along the hepatocyte trajectory from E13.5 no longer express the epitope for the anti-Liv2 antibody used during FACS, and thus were not subject to sequencing.

Together, our scRNAseq analysis suggested that the E10.5 embryonic liver harbours distinct sub-populations of liver progenitors that co-exist within the hepatoblast pool; a Lgr5+ sub-population that contributes to both hepatocyte and cholangiocytes and a previously unrecognized sub-population of already cholangiocyte committed cells that has already downregulated *Lgr5* and started its specification to the cholangiocyte fate.

### Lgr5 marks the apex cells within a E9.5 heterogenous hepatoblast pool

Our lineage tracing and single cell RNA sequencing data showed Lgr5 labels bi-potential hepatoblasts which differentiate towards hepatocyte or cholangiocyte fates. This is indicative of a hepatoblast hierarchy, and suggested Lgr5 as a potential marker of its apex. Quantifying the number of tracing events as well as their contribution to the postnatal tissue provides information on the potency and commitment of a given population in the developing tissue. To determine whether Lgr5+ cells reside at the apex of a developmental hierarchy, we reasoned that the cell composition of their clonal progeny must reflect quantitatively the corresponding proportions in tissue. Therefore, we quantified the proportion of labelled hepatocytes and cholangiocytes following lineage tracing from *Lgr5-CreERT2* at E9.5 (Fig. 6A) and compared the proportions to the homeostatic distributions (Fig. 6B). We found that the homeostatic proportion of hepatocytes and cholangiocytes in the mouse postnatal liver is 96.6 ± 0.6% and 3.4 ± 0.6%, respectively, (Fig. S6); in agreement with previous reports in rats (Blouin et al., 1977). Remarkably, we found that lineage tracing with *Lgr5-CreERT2* at E9.5 resulted in labelled cells in which 96.7 ± 0.5% were hepatocytes and 3.3 ± 0.5% were cholangiocytes, the same proportion of labelled hepatocytes and cholangiocytes as the homeostatic liver (Fig. 6C). Therefore, although we cannot rigorously rule out the possible contribution of a parallel Lgr5-cell lineage that produces differentiated progeny in tissue proportions, these results strongly suggest that Lgr5 expression marks hepatoblasts that constitute the apex of the differential hierarchy.

**Figure 6:**
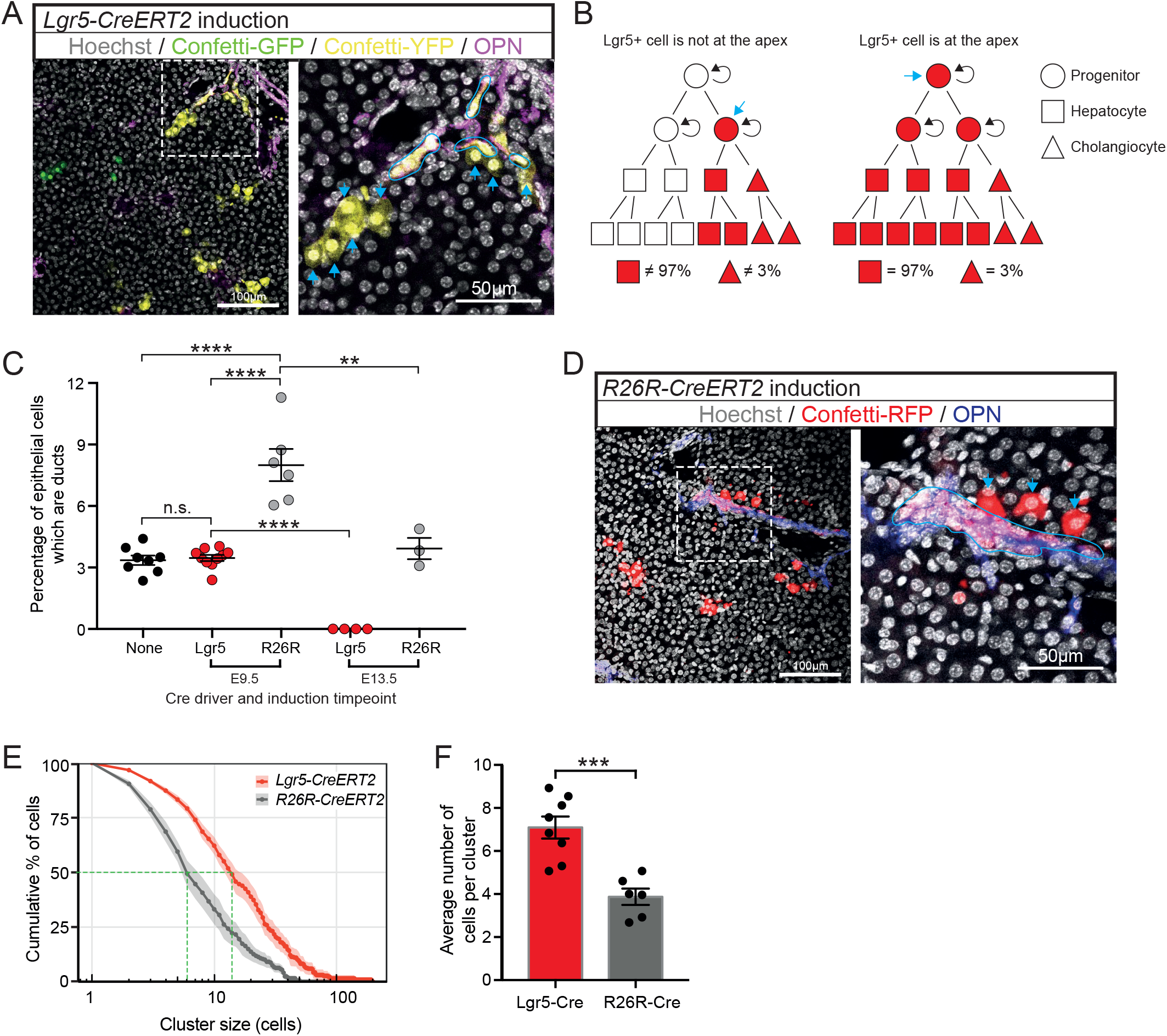
Lgr5+ hepatoblasts are at the apex of the hepatoblast hierarchy. (A-E) Pregnant females from both, the Lgr5-Cre (Lgr5-CreERT2) and the ubiquitous Cre (Rosa-CreERT2, R26RCre) were injected at E9.5 or E13.5 of gestation in order to lineage trace the early hepatoblast pool. Livers were collected postnatally and assessed for the contribution of the labelled descendants to the ductal and hepatocyte pools on the postnatal liver. Clone composition, clone number and clone size were quantified. (A) Representative images of Confetti-labelled descendants following tamoxifen induction to *Lgr5-CreERT2; R26Rconfetti* mouse at E9.5 and liver collection at P17. Magnified area presents ductal cells (Osteopontin+, purple) outlined in blue. (B) Schematic displaying the possible outcomes of labelling proportions following lineage tracing of Lgr5+ cells at E9.5 depending on where Lgr5+ cells are in the hepatoblast hierarchy (indicated with the blue arrow). If E9.5 Lgr5+ hepatoblasts are at the apex of their hepatoblast hierarchy, it is expected that their contribution to the postnatal liver should represent the homeostatic proportions of hepatocytes and ductal cells in the liver (97% vs 3%) as detailed in Blouin et al 1977 and in supplementary figure S6D. In the left panel, Lgr5 (blue arrow) is not at the apex, hence the homeostatic proportion is not achieved. On the contrary, in the right panel, Lgr5+ cells are at the apex, and that generates the homeostatic proportions. (C) Graph shows that 3.5% ± 0.5% of labelled epithelial cells following induction from *Lgr5-creERT2* were ductal, which is equivalent to the homeostatic percentage of ductal cells in the postnatal liver (none Cre driver). In contrast, the percentage of labelled ductal cells using *R26R-CreERT2* at E9.5 was significantly higher. At E13.5, lineage tracing from the *Lgr5-CreERT2* allele resulted in no labelled ductal cells, whilst induction from the allele at E13.5 gave the same homeostatic proportion (mean±SEM, each data point represents an individual liver). Analysis of postnatal livers was conducted at timepoints P0-P30; later time points were not considered to prevent homeostatic cellular turnover confounding the data. **, p<0.01; ***, p<0.001. (D) Representative images of Confetti-labelled descendants following tamoxifen induction to *R26R-CreERT2;R26Rconfetti* mouse at E9.5 and with liver collection at P14. (E) Cumulative distribution of cluster size frequency at P14-P30, comparing labelled clusters derived from Lgr5+ cells (Lgr5-creERT2) and the bulk population (R26R-CreERT2) induced at E9.5 (mean±SEM, n≥6). Tracing from Lgr5+ cells results in larger clusters than tracing from the bulk population (F), suggesting that Lgr5+ cells have greater proliferative potential than the bulk population at E9.5 (mean±SEM).

When analysing our single cell RNA sequencing data, we also found that at E10.5 there were cholangiocyte-like cells that did not express Lgr5, suggesting that these were cholangiocyte-committed hepatoblasts even at this very early time point. To formally investigate whether this was a genuine functional heterogeneity or was only reflecting transcriptional heterogeneity at this time point we turned to a second lineage tracing strategy using an ubiquitous and unbiased driver: the *R26R-CreERT2*. Lineage tracing from *R26R-CreERT2* will label all cell types in the developing liver, including Lgr5+ and Lgr5-hepatoblasts, and therefore labelled hepatocytes and cholangiocytes in postnatal livers will represent descendants of any hepatoblasts labelled at E9.5 (Fig. 6D). Strikingly, we found a significantly higher proportion of labelled cholangiocytes compared to the homeostatic proportion when labelled with the unbiased *R26R-CreERT2* allele at E9.5 *(R26R-CreERT,* 7.7 ± 1.9% cholangiocytes *vs* 3.3 ± 0.5% in homeostasis and with *Lgr5-CreERT2)* (Fig. 6C). These results were confirmed using multiple multicolour R26R-reporter alleles (R26R-Confetti and R26R-Rainbow). These findings are in agreement with our scRNAseq data, in which we had observed that E10.5 hepatoblasts were already committed to a cholangiocyte fate.

Interestingly, and in contrast to induction at E9.5, Cre induction from the *R26R-CreERT2* allele at E13.5 gave rise to labelled hepatocytes and cholangiocytes in homeostatic proportions (Fig. 6C), whilst lineage tracing from the *Lgr5-CreERT2* allele at E13.5 gave rise solely to labelled hepatocytes (Fig. 6C), suggesting that Lgr5+ cells lose their potency and position in the hierarchy during the developmental progression. These results indicate that hepatoblasts are not only heterogeneous in progenitor potential but their competence to generate hepatocytes and cholangiocytes changes between E9.5 and E13.5.

In addition to the identity of labelled cells, the size of labelled clusters generated from the *Lgr5-CreERT2* and *R26R-CreERT2* alleles was quantified as a proxy for the proliferative potential of the initially labelled hepatoblast. We found that tracing with the *Lgr5-CreERT2* allele at E9.5 gave rise to larger clusters of labelled cells than tracing with the *R26R-CreERT2* allele (Fig. 6E,F). The larger cluster sizes from Lgr5+ hepatoblasts indicate that these cells have a greater proliferative potential than the bulk hepatoblast population, again suggestive of their position at the apex of the hepatoblast hierarchy.

Altogether combined, these results lead us to conclude that the E9.5 hepatoblast population is indeed functionally heterogeneous, with Lgr5+ hepatoblasts residing at the apex of the E9.5 hierarchy and a population of non-Lgr5+ hepatoblasts exhibiting a previous unidentified early commitment to the cholangiocyte fate.

## Discussion

The Wnt target gene *Lgr5* (leucine-rich-repeat-containing G-protein-coupled receptor 5) has been described as a marker of stem cells in non-damaged, self-renewing tissues, such as the intestine, stomach and hair follicles, as reviewed in (Barker et al., 2010). In the adult liver, Lgr5 is lowly expressed during homeostasis. However, upon damage Lgr5 becomes highly upregulated in a subset of cells, which contribute to the regeneration of both, hepatocytes and ductal cells. Similarly, Lgr5 is also upregulated in homeostatic liver ductal cells, when cultured as self-renewing bi-potential liver organoids (Huch et al., 2013b). Here we found that Lgr5 marks a previously unknown bi-potent Lgr5+ population, which resides at the apex of an E9.5 heterogenous hepatoblast pool.

To date, bi-potentiality of hepatoblasts has only been shown at the population level (Yanagida et al., 2016), however, at least *in vivo,* there has been no experimental proof regarding bi-potentiality of individual hepatoblast cells. A recent report showed that a labelled Foxa2+ definitive endoderm cell induced at E7.75 gives rise to cells moving towards hepatocyte and cholangiocyte fates at E16.5, suggesting that at least, before hepatic specification at E7.75, the definitive endoderm progenitors are multipotent (El Sebae et al., 2018). Similarly, *in vitro,* Dlk+ embryonic liver cells at E14.5 were found to express markers of both hepatocyte and cholangiocyte lineages (Tanimizu et al., 2003), again suggestive of the bi-potent nature of hepatoblast cells. However, formal proof of *in vivo* bi-potential hepatoblasts has not yet been provided. Here, using lineage tracing with a multicolour reporter we unequivocally demonstrate that E9.5 Lgr5+ hepatoblasts are indeed bi-potential *in vivo,* as single clones consisting of cholangiocytes and hepatocytes are present at the portal triad (Fig. 3C,D). It should be noted, though, that we also found clones formed of hepatocytes without cholangiocytes throughout the liver; including in the portal region (zone 1). These can have two possible explanations: either a subset of Lgr5+ cells is unipotent for hepatocyte fate and others are bi-potent, or alternatively, all the Lgr5+ cells are bi-potent. The first option (only a subset is bi-potent) implies that there is heterogeneity within the Lgr5+ population regarding their potentiality. In that regard, our single cell data, where we find Lgr5+ cells in both the hepatoblast and the hepatocyte clusters, suggest that this could indeed be a plausible scenario. Alternatively, one could hypothesise that all the Lgr5+ cells are bi-potent but depending on the external signals received according to the specific position of the original Lgr5+ progenitor cell, they can differentiate into one or two cell types. This implies that developmental stage and local environment could be critical in defining the final fate of a given Lgr5+ hepatoblast. In that regard, the fact that embryonic Lgr5+ cells isolated by FACS and cultured *in vitro* were sensitive to the growth factors present in the culture medium and committed either to the cholangiocyte or hepatocyte lineage according to media composition would argue in support of this latter argument (Fig. 4). It is tempting to speculate that by retaining Lgr5+ cells at defined positions during liver growth by maintenance of a specific local environment, differentiation into cholangiocytes would occur as daughter cells exited such a niche. This is consistent with current evidence of the discontinuous growth of the liver ductal network as reviewed in (Ober and Lemaigre, 2018), although there is at present no direct evidence for a role of Lgr5+ cells in directing liver morphogenesis.

Interestingly, lineage tracing at E13.5 from the *Lgr5-CreERT2* allele resulted in labelling of only hepatocytes (Fig. 6C,S1B), while E13.5 lineage tracing from the *R26R-CreERT2* allele resulted in labelling of both, hepatocytes and ductal cells, in the homeostatic proportions (96.4% and 3.6%, respectively). These results underline the continual shift in cell potency and cell surface marker expression throughout development of the liver and are consistent with other reports in which a single cell surface marker is not adequate to define a particular cell type (hepatoblast, hepatocyte or cholangiocyte) throughout the entirety of liver development (Tanaka et al., 2009). Instead, a set of two or more cell surface markers will have to be used to define each cell type at specific stages of development. In that regard we found that while Liv2 is indeed a good marker of the hepatoblast pool at E10.5, it is not appropriate to identify unbiased hepatoblasts at E13.5, as it seems to mark hepatoblasts already biased towards the ductal fate.

In contrast to the widely accepted view that differentiation of hepatoblasts into cholangiocytes occurs from E13.5 onwards (Gordillo et al., 2015), our results provide the functional demonstration that heterogeneity indeed already exists at E9.5 of liver development. Our single cell RNA sequencing data shows that even as early as E10.5 there is heterogeneity within the hepatoblast population, with some cells already moving towards cholangiocyte or hepatocyte fates. We identify sub-populations of hepatoblasts that express Lgr5 whilst other sub-populations do not. Further, some of these Lgr5-cells already express markers of the cholangiocyte fate. In support of the scRNAseq data, our functional studies which fate map E9.5 liver progenitors using lineage tracing from either *Lgr5-CreERT2* or *R26R-CreERT2* demonstrate the existence of both Lgr5+ and Lgr5-hepatoblasts already at E9.5 (Fig. 6C). Importantly, induction of lineage tracing at E9.5 using *Lgr5-CreERT2,* but not *R26R-CreERT2,* resulted in labelled postnatal hepatocytes and cholangiocytes in homeostatic proportions (97% hepatocytes vs 3% cholangiocytes), implying that Lgr5+ cells functionally behave as a genuine bi-potent hepatoblast and are indeed at the apex of its hepatoblast hierarchy. On the contrary, the unbiased *R26R-CreERT2,* gave rise to a higher proportion of cholangiocytes at E9.5, compared to the homeostatic or Lgr5+ descendants, arguing in favour of an already cholangiocyte-committed hepatoblast sub-population, negative for Lgr5, in the E9.5 developing liver. This result suggests that E9.5 Lgr5+ cells are at the apex of their hierarchy, i.e., are bi-potent and equipotent, and are able to give rise to already lineage-restricted ductal progenitors that downregulate Lgr5 and expand in order to contribute to the postnatal ductal pool. While our results demonstrate that Lgr5 expression overlaps with the apex of an hepatoblast pool, they do not show functionally that Lgr5 expression defines the apex of the hepatoblast pool. Lgr5 expression could be subjected to local environmental factors and just mark a subpopulation of cells that receives high Wnt signalling. Then, one could speculate that Lgr5 negative hepatoblasts with the very same potency as the Lgr5+ ones also reside at the apex of their own hierarchies. This implies that populations of equally potent hepatoblasts, some of which express Lgr5 and are bi-potent, co-exist at this time point in development. If these additional hepatoblasts populations indeed exist, what is their identity and bi-potentiality are still questions that remain to be addressed.

In summary, here, using a combination of lineage tracing, organoid cultures and scRNAseq analysis we describe for the first time that the E9.5 hepatoblast pool is heterogeneous, not only at the RNA, but also at the functional level. Within the different E9.5 hepatoblast sub-populations, we find that Lgr5 marks a previously unknown bi-potent Lgr5+ population, which resides at the apex of its E9.5 hepatoblast hierarchy. Furthermore, we also describe a previously un-identified sub-population of already cholangiocyte-committed cells that do not express Lgr5. To our knowledge, this is the first report that recognizes the functional heterogeneity of the E9.5 hepatoblast pool and the first demonstration that *Lgr5* is a *bona-fide* marker of early bi-potent hepatoblasts in the developing liver. Our studies raise further questions about the nature of Lgr5 in liver development and liver morphogenesis. Wnt signalling has been implicated in liver growth (McLin et al., 2007; Micsenyi et al., 2004; Tan et al., 2008), however its role in determining the potency of hepatoblasts is unknown. Elucidating the functional role, if any, for Lgr5 in liver development could help to clarify the part played by Wnt signalling. In our hands, KO of the Lgr5 gene did not result in an apparent liver phenotype (Fig. S7). However, the presence of other homologues, like Lgr4, which is expressed during liver development (Camp et al., 2017), could compensate for the loss of function of Lgr5, as it is reported for the adult intestine (de Lau et al., 2011). Therefore, it remains unclear whether Lgr5 *per se* has a functional role in liver development. Similarly, cell ablation studies would be required to address whether the Lgr5+ hepatoblast rather than the Lgr5 gene *per se* is indeed required during development. Due to the wide-spread expression of Lgr5+ stem cells in the adult and in other embryonic tissues, it is not trivial to assess the functionality of Lgr5+ cells in a specific tissue. We have shown that isolated Lgr5+ hepatoblasts can be cultured *in vitro,* and so this may provide a reductionist system in which we can test the requirement for Lgr5 in establishing or maintaining bi-potency in hepatoblasts without the confounding effects of signalling from other tissues. Future studies would aim at addressing these questions.

## Materials and Methods

### Mouse strains and animal work

Lgr5-CreERT2 (Huch et al., 2013b), Lgr5-GFP (Barker et al., 2007), R26R-TdTomato (Madisen et al., 2010), R26R-Confetti (Snippert et al., 2010) and R26R-CreERT2 (Ventura et al., 2007) and R26R-Rainbow1.0 (Livet et al., 2007) mice were described previously. All mouse experiments have been regulated under the Animals (Scientific Procedures) Act 1986 Amendment Regulations 2012 following ethical review by the University of Cambridge Animal Welfare and Ethical Review Body (AWERB) and have been performed in accordance with the Home Office license awarded to M.H.

### Tamoxifen induction

Lineage tracing was performed using the R26R-TdTomato or R26R-Confetti or R26R-rainbow reporter in combination with a temporally inducible Cre, either Lgr5-CreERT2 or R26R-CreERT2. To induce Cre activity tamoxifen (Sigma, T5648) was administered by intraperitoneal injection of the pregnant female at E9.5. Tamoxifen doses were dependent on the reporter line and Cre line used. For details refer to Supplementary dataset 2. Embryos and pups (male and females) were then collected at specified time points according to the experiment.

### Tissue preparation and immunostaining

Embryonic and postnatal livers were dissected and fixed for 2 h or 24 h, respectively in 10% neutral-buffered formalin (Sigma-Aldrich) at 4°C. Postnatal livers were embedded in 4% low melting point agarose (BioRad Laboratories) and sectioned at 100μm using a Leica VT1000S microtome. To reduce nonspecific staining and permeabilize the sample, samples were incubated with a 2% donkey serum, 1% Triton, 5% DMSO in PBS solution overnight at 4°C. Primary antibodies were then applied at appropriate dilutions for 48 h at 4°C. Samples were washed and secondary antibodies applied at dilution 1:250 for 48 h at 4°C. Nuclei were counterstained with Hoescht 33342 (1:1000, Invitrogen) for 30 min at room temperature.

### Confocal Imaging

Samples were imaged on a SP8 White Light inverted confocal microscope (Leica Microsystems) through a 10x or 20x objective using a Leica application suite X Software. Optical sections were acquired at 2μm intervals. Images were processed using Fiji.

### Isolation of cells for single cell RNA sequencing and *in vitro* culture

Lgr5GFP^het^ and Lgr5GFP^−/-^ embryos were collected at the specified timepoints and screened for GFP signal under an epi-florescence microscope. Once classified according to phenotype, livers were collected and minced before enzymatic digestion. Enzymatic digestion was performed at 37°C with Wash medium (constituting DMEM+ GlutaMAX (Invitrogen) supplemented with 1% FBS and 1x penicillin/streptomycin) containing 0.125 mg/ml Collagenase Type I (Sigma-Aldrich) and Dispase II (Gibco) and 0.1 mg/ml DNase (Sigma Aldrich). The incubation time for enzymatic digestion was approximately 40 min for E10.5 livers and 2 h for E13.5 livers. Once the digestion to single cells was confirmed by visual inspection, samples were filtered through a 40 μm pore size nylon cell strainer (Falcon) and centrifuged at 400 g for 5mins. The pellet was resuspended in blocking solution (Wash medium with 2% FBS, Rho kinase inhibitor Y27632 (Sigma Aldrich) and 0.1 mg/ml DNase for 20 min. Cells were then centrifuged at 400 g for 5 min and incubated with primary antibody against Liv2 (1:100, MBL) in washing medium supplemented with rock inhibitor and DNase for 40 min on ice. Cells were then pelleted at 400 g for 5 min and washed. Cells were incubated with APC anti-rat (Biolegend) for anti-Liv2, CD31-PE/Cy7 (abcam) and CD45-PE/Cy7 (Bioscience) diluted in washing medium for 40 min on ice. The sorting strategy consisted of a population of single cells that were sequentially gated based on cell size (forward scatter, FSC, versus side scatter, SSC), singlets (pulse width vs FSC) and Liv2-APC positivity. Finally, CD45-PE/Cy7 (BD Biosciences), CD31-PE/Cy7 (Abcam) antibodies were used in order to exclude blood cells and endothelium. Liv2^+^/CD31^−^/CD45^−^ (bulk hepatoblast pool) or Liv2^+^/CD31^−^CD45^−^GFP^+^ cells (named as Lgr5^+^ cells) were used for further analysis.

For single cell RNA sequencing experiments cells were sorted on an influx Cell Sorter (BD Biosciences). Single cells were collected in non-skirted PCR plates containing lysis buffer (0.2% triton (Sigma triton X-100 solution) in 1 U per μl RNase inhibitor (Thermo Fisher Scientific) in DEPC-water (Ambion)). Plates were then vortexed and centrifuged at 2000 rpm for 2 mins and kept at −80°C. For 3D *in vitro* culture cells were sorted on a MoFlo into sort medium (Advanced DMEM/F12 (GIBCO) supplemented with 1% penicillin/streptomycin, 1% Glutamax, 10 mM HEPES, 1x B27 supplement (without vitamin A), 1.25 mM N-acetyl-l-cysteine, 10% (vol/vol) Rspo-1 conditioned medium, 10 mM Nicotinamide, 10 nM recombinant human (Leu15)-gastrin I, 50 ng/ml recombinant mouse EGF, 100 ng/ml recombinant human FGF10, 25 ng/ml recombinant human HGF, 5 μM A8301, and 10 μM Y27632.

### Single cell RNA sequencing

scRNA-seq sample preparation was performed with an adapted version of Smartseq2 (Picelli et al., 2014). cDNA was reverse transcribed using 50 U. reaction SmartScribe Reverse Transcriptase (Takara ClonTech) without Betaine and MgCl_2_ and amplified using KAPA HiFi Hotstart polymerase (Roche). Illumina Nextera XT DNA preparation kit was used to prepare libraries and pooled libraries were sequenced using the Illumina HiSeq 4000 system (single-end 50 bp reads). The quality of the reads was examined with FastQC (http://www.bioinformatics.babraham.ac.uk/projects/fastqc/). The reads were aligned to genome version GRCm38, with the 92 Spike-in transcript sequences added, using STAR (v2.6.0c) and Ensemble gene annotation version 93 (Kersey et al., 2018). subread (v1.6.2) was used to count uniquely aligned reads using the same Ensemble annotation and to create the count matrix. Further analysis was performed using scanpy (v1.3.3) (Wolf et al., 2018b). For quality control of cells, the following quality metrics were calculated for each cell: (1) the percentage mitochondrial transcript reads, (2) the percentage of Spike-In reads, (3) the total number of reads, and (4) the log10 transformed number of genes with at least one read. Only cells with (1) less than 20% of mitochondrial reads, (2) less than 25% Spike-In reads and (3) more than 100.000 reads were considered for downstream analysis. As the log10 transformed number of genes with at least one read (4) showed clear batch effects, the four different thresholds 3.6, 3.5, 3.7, and 3.5 were applied to the four different sorts and only cells exceeding these thresholds passed quality control. In total, 653 (69%) of 943 cells were considered for downstream analysis. Because an initial principal component analysis revealed batch effects between the biological replicates from experiments 1, 2 and 4 (group 1) on the one hand and experiment 3 (group 2) on the other hand, batch correction between those two groups was performed: For each group, only genes expressed in at least 3 cells were considered. The counts in each group were normalised using size factors computed with the scran (v1.8.4) function computeSumFactors (parameters: min_mean=1.0, size=seq(20, 100, 5)) (Lun et al., 2016). For each group, highly variable genes were detected using the scanpy function filter_genes_dispersion (parameter: max_mean=8) and the intersection of both gene sets, which contained 1766 genes, was used for further analysis. Batch effects between the two datasets were corrected by matching mutual nearest neighbours in the implementation of mnnpy (v0.1.9.3) (parameters: svd_mode=‘irlb’) (Lun et al., 2016). On the resulting count matrix, a principal component analysis was performed. t-SNE dimensionality reduction was performed on the first 20 principal components using the MulticoreTSNE implementation (paramters: perplexity=80, early_exageration=12) (Amir et al., 2013). To perform Louvain clustering, the 15-nearest neighbours graph was computed on the first 20 principal components. Using Louvain clustering with the resolution parameter set to 0.05, 3 clusters were obtained (Levine et al., 2015; Subelj and Bajec, 2011). Differentially expressed genes were detected by performing a Wilcoxon rank-sum test on the raw counts comparing each cluster against the union of the other two clusters as implemented in scanpy’s rank_genes_groups function. To define marker genes for the clusters at specific embryonic stages, the cells were grouped according to cluster and stage and a Wilcoxon rank-sum test was performed as described above. For the sub-clustering of the cholangiocyte-like cluster, a principal component analysis was performed on those cells and then clustering was performed as above with the resolution parameter set to 0.5. Differentially expressed genes between the two resulting sub-clusters were detected as described above. The diffusion maps were calculated using the scanpy function diffmap with the width of the Gaussian connectivity kernel being implicitly determined by the distance to the 100 nearest neighbours in the space of the 20 first principal components (Coifman et al., 2005; Haghverdi et al., 2015). Diffusion pseudotime was calculated using scanpy’s dpt function using the cell with minimal diffusion component 1 as root cell (Haghverdi et al., 2016; Wolf et al., 2018a).Cell cycle phases were assigned using cyclone and the pre-trained mouse cycle markers contained in the scran package (Scialdone et al., 2015). Cells were classified as Lgr5 positive on the transcript level if they had more than 10 reads of Lgr5.

### 3D culture of embryonic liver cells

Following isolation as described in isolation section above, the cells were pelleted at 400g and seeded in Matrigel (BD Biosciences) and cultured either with the ‘hep’ protocol or ‘ductal’ protocol. The ‘hep’ method involves culturing for the first 3 days in Advanced DMEM F12 supplemented with Penicillin/Streptomycin, Glutamax and HEPES (Gibco), 1xB27 (Gibco), 500 nM n-Acetylcysteine (Sigma), 10 mM Nicotinamide (Sigma), 100 ng/ml FGF10 (Peprotech), 100 ng/ml FGF7 (Peprotech), 50 ng/ml HGF (Peprotech), 10 nM Gastrin (Sigma), 50 ng/ml EGF (Peprotech), 1 nM A83-01 (Tocris Bioscience), 3μM CHIR 99021 (Tocris Bioscience), 15% R-spondin 1 conditioned medium (in house), and 10 μM Rock inhibitor Y-27632 (Sigma). From day 3 onwards the culture medium was modified by the exclusion of FGF7. The ‘ductal’ method consists of culturing for the first 3 days in Advanced DMEM F12 supplemented with Penicillin/Streptomycin, Glutamax and HEPES 1xB27, 500 nM n-Acetylcysteine, 10 mM Nicotinamide, 100 ng/ml FGF10, 50 ng/ml HGF, 10 nM Gastrin, 25 ng/ml Noggin (Peprotech), 50 ng/ml EGF, 1 nM A83-01, 10μM Forskolin (Tocris Bioscience), 10% R-spondin 1 conditioned medium, 30% Wnt conditioned medium (in house) and 10 μM Rock inhibitor Y-27632 (Sigma). From day 3 onwards the culture medium was modified by the exclusion of the Wnt conditioned medium and removal of Noggin. After several days in culture, organoid structures with either a cystic (duct) or solid (hep) form arose. Cultures were split at 1:2 ratio after 14-20 days.

## Acknowledgements

We would like to acknowledge the Gurdon Institute Animal facility and Richard Butler at the Gurdon Institute Imaging Facility for microscopy and image analysis support. Andy Riddell and Simon McCallum for cell sorting. Robert Arnes-Benito for help in sectioning in the early phases of the project. Rachel Tan for help with counting.

## Author contributions

Conceptualization: N.P., C.J.H, B.D.S, M.H.; Methodology: N.P., C.J.H., W.L., M.H.; Software: F.R., S.R.; Validation: N.P., E.M.E.; Formal analysis: N.P., F.R., S.R.; Investigation: N.P., C.J.H., E.M.E.; Resources: B.D.S., M.H.; Data curation: F.R., S.R.; Writing - original draft: N.P., M.H.; Writing - review & editing: N.P., C.J.H., F.R., S.R., B.D.S., M.H.; Supervision: B.G., S.R., B.D.S., M.H.; Project administration: M.H.; Funding acquisition: B.D.S., M.H.

## Data availability

Generated datasets have been deposited in GEO under accession number GSE123103.

---

**Supplementary Fig. 1:**
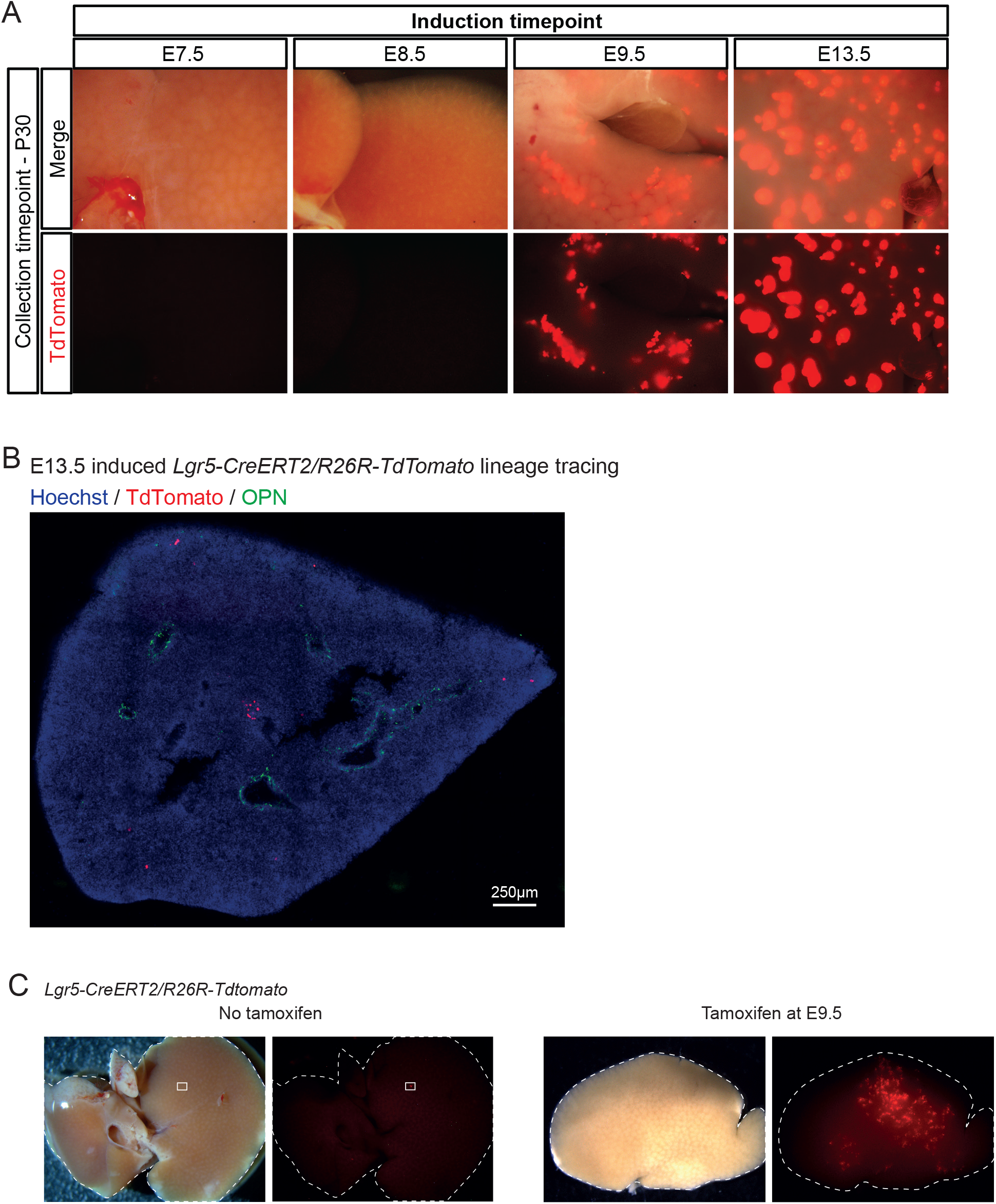
Lgr5 labels hepatoblasts from E9.5 to at least E13.5 of liver development. (A-C) *Lgr5-ires-CreERT2^hom^; ROSA-TdTomato^hom^* males were breed with MF1-WT females in order to generate the compound mice *Lgr5-ires-CreERT2*+/-;*R26R-TdTomato*+/-. (A) Cre activity was induced in *Lgr5-ires-creERT2*+/-;*R26R-TdTomato*+/- embryos at the indicated timepoints and livers collected at P30. Expression of TdTomato was detected in postnatal livers only if induction was at E9.5 or later, suggesting that Lgr5 is expressed in the developing liver from E9.5 onwards. Scale bar, 2mm. (B) Lineage tracing from *Lgr5-ires-creERT2*+/-; *R26R-TdTomato*+/- embryos induced at E13.5 resulted in only hepatocyte progeny. (C) TdTomato tracing was almost never detected in non-tamoxifen induced controls. Only in one mouse we detected <100 cells labeled in the liver. In contrast, the tracing events in the livers of the tamoxifen induced embryos was clear with several thousands of cells labeled. Both the non-induced and induced livers were collected at P14. This minute number of labelled cells in the non-induced livers does not affect our interpretations in the lineage tracing experiments.

**Supplementary Fig. 2:**
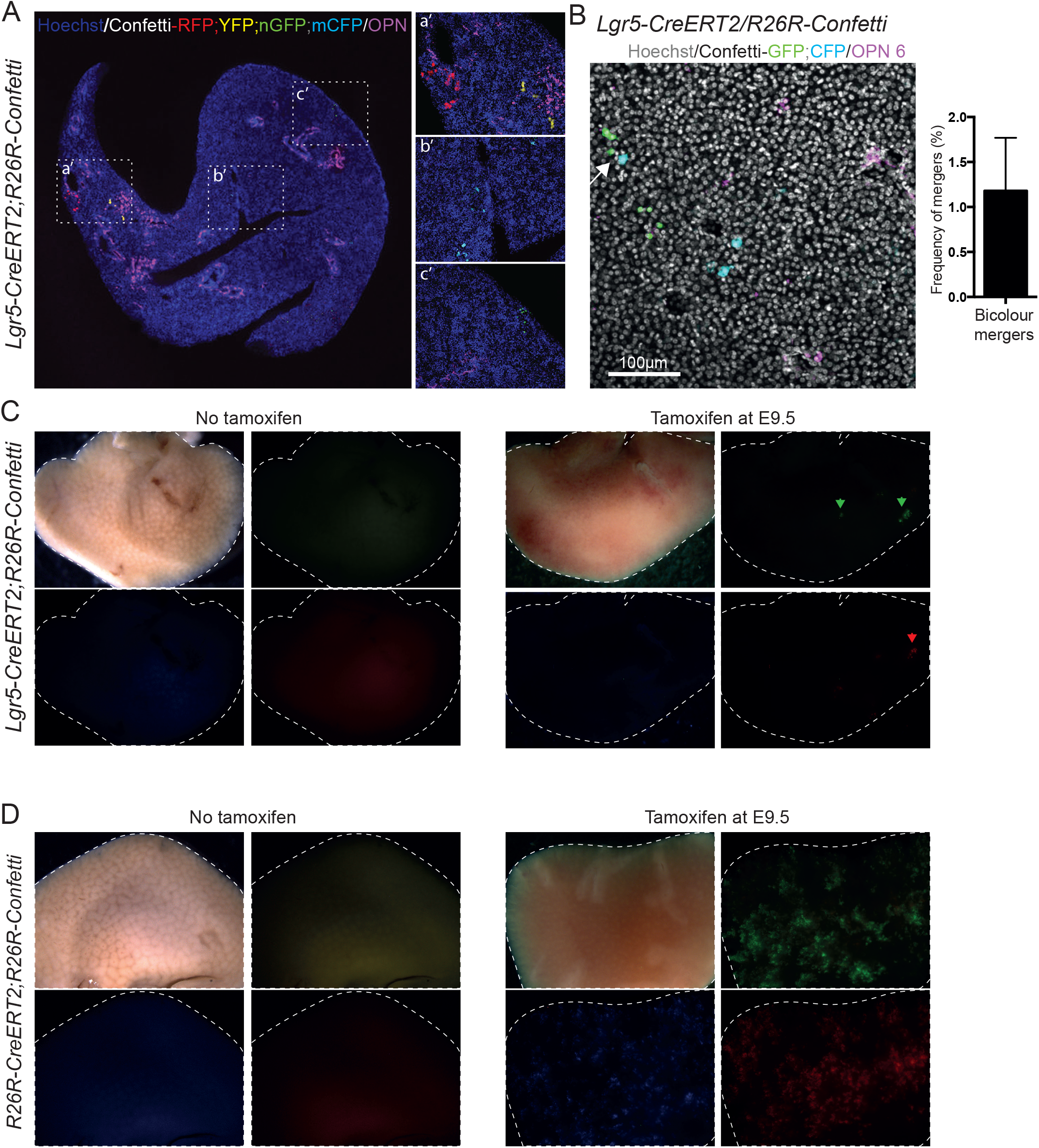
Lineage tracing from Lgr5-ires-creERT2;R26R-confetti and R26RCre;R26R-confetti mice induced at E9.5. (A-C) *Lgr5-ires-CreERT2;R26R-Confetti* mice were generated by breeding the Lgr5-ires-CreERT2hom with the multicolour Confetti reporter R26R-Confettihom and liver tissues were collected postnatally. (A) Labeling of the E9.5 embryonic livers of *Lgr5-ires-CreERT2;R26R-Confetti* mice results in cells labelled in one of four colours (RFP, YFP, mCFP, nGFP). A representative image of a liver section presenting clones in each of the 4 colours is shown. Tissue was co-stained with Osteopontin to visualize the ductal cells (OPN, magenta). Nuclei were counterstained with Hoecsht. a’, magnified area showing a red and a yellow clone. b’, magnified area showing a mCFP clone. c’, magnified area showing a nGFP clone. Note that no merging of clones is observed. (B) Bicolour-merging events, i.e., clones of different colours merging, were rarely detected. Representative image of one of the 2 merging events observed is shown. Example of a merging event between a CFP and GFP clone following induction at E9.5 in *Lgr5-ires-creERT2*+/-;*R26R-Confetti*+/- embryos. Scale bar, 100μm. Graph represents the proportion of bicolour-merging events identified in the 3 livers analysed (1.2% ± 1.0). Results represent mean ± SEM of the merging events found in the n=3 livers analysed (n=1, liver_1; n=1, liver_2, n=0, liver_3). (C-D) Tracing events from the R26R-confetti reporter are only in combination with the *Lgr5-CreERT2* (C) or *R26R-CreERT2* (D) driver are only detected upon tamoxifen administration and never found in non-induced mice.

**Supplementary figure 3:**
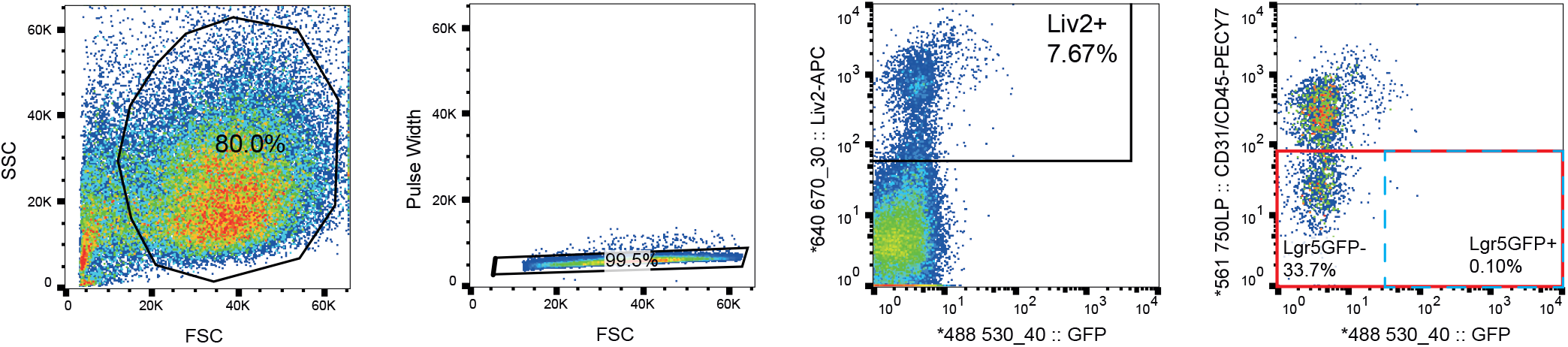
Lgr5+ hepatoblast sorting strategy. *Lgr5-EGFP-IRES-CreERT2^het^* mice were breed and embryos collected at E10.5 of gestation. *WT* and *Lgr5-EGFP-IRES-CreERT2^het^* littermate embryos were scored for the presence of eGFP in the cranial area. Then, embryos were split by genotype and liver tissues collected and processed for cell isolation and single cell dissociation. Cells were stained with the hepatoblast marker Liv2, the endothelial maker CD31 and pan-haemopoietic marker CD45 as described in methods. Sorted cells were obtained following a sequential gating strategy where cells were first gated by FSC vs SSC, then FSC vs Pulse with was used to identify singlets and then gated for Liv2+ (bulk hepatoblasts, Liv2+CD31-CD45-) or Liv2+eGFP+ (Lgr5+ hepatoblasts, (Liv2+CD31-CD45-GFP+ (blue dashed box)).

**Supplementary Figure 4:**
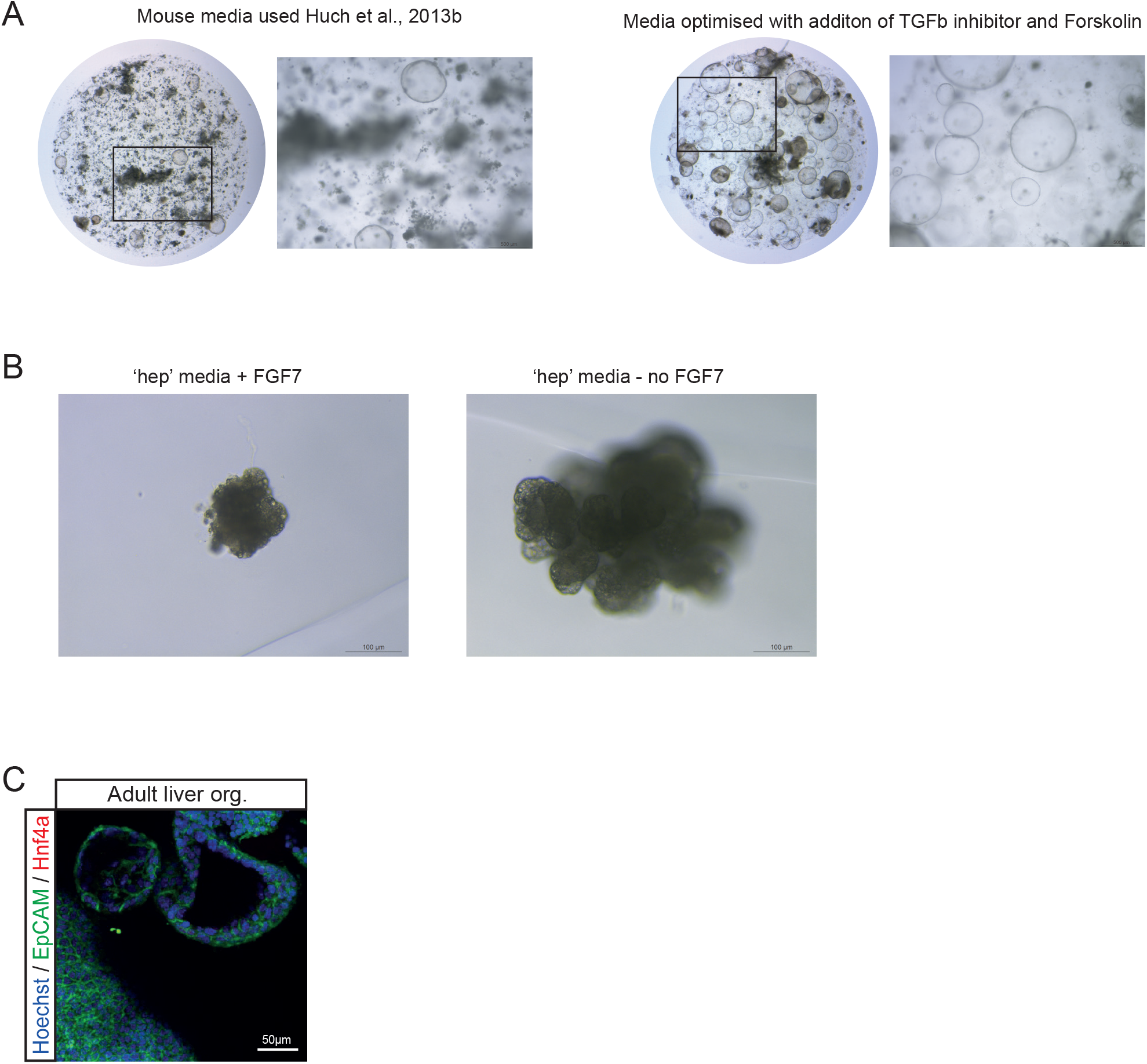
Generation of mouse embryonic liver organoids. (A-B) Embryonic liver organoids were generated from bulk hepatoblasts obtained from E11.5 *WT* or *Lgr5-EGFP-IRES-creERT2* embryos dissociated as described in methods. (A) Isolated cells were embedded in Matrigel and cultured in our mouse liver organoid medium (Huch et al., Nature 2013) containing EGF, Noggin, Rspondin1, FGF10, HGF, Nicotinamide (left panel) or in the same medium supplemented with 1nM A8301 and 10uM Forskolin (optimized duct medium, right panel) as described in methods. This resulted in a evident increase in the number and size of organoids formed. (B) Isolated cells embedded in Matrigel were also cultured in the recently published hepatocyte (hep) medium that sustains human embryonic liver growth in vitro (ref Hu et al., 2018). Removal of FGF7 resulted in a significant improvement on the expansion of the organoids. For details refer to methods. (C) Immunofluorescence staining for the ductal marker EpCAM and the hepatocyte marker HNF4 in adult liver organoids cultured in our standard organoid medium as described in (Huch et al., Nature 2013). These served as positive control for the stainings shown in Figure 4.

**Supplementary Figure 5:**
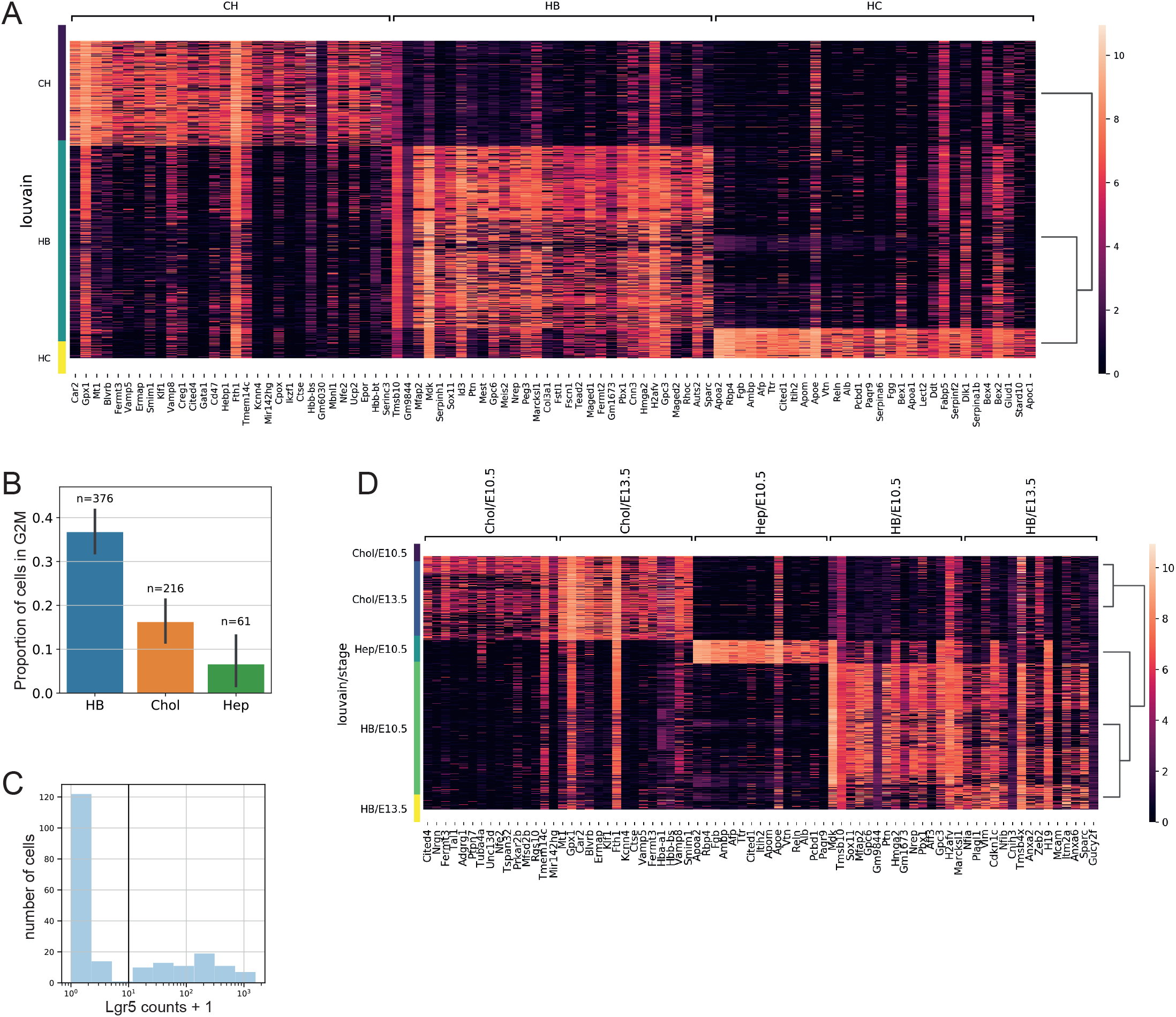
scRNAseq of hepatoblasts reveals heterogeneity in the hepatoblast population. (A) Heatmap displaying the 30 most differentially expressed genes in each cluster as detailed in Supplementary Dataset 1. (B) Proportion of cells in G2M phase of the cell cycle in the hepatoblast cluster (HB), cholangiocyte-like cluster (Chol) and hepatocyte-like cluster (Hep) (mean±95% confidence intervals). (C) Histogram of the distribution of Lgr5 counts shows a bimodal distribution. A threshold of 10 counts is used to define a cell as Lgr5+ on the transcript level. Using this threshold, we find that 2% of the bulk cells at E10.5 are Lgr5+ at the transcript level. (D) Heatmap displaying the differentially expressed genes by time point. For extended list see Supplementary Dataset 1_S2-S6.

**Supplementary Figure 6:**
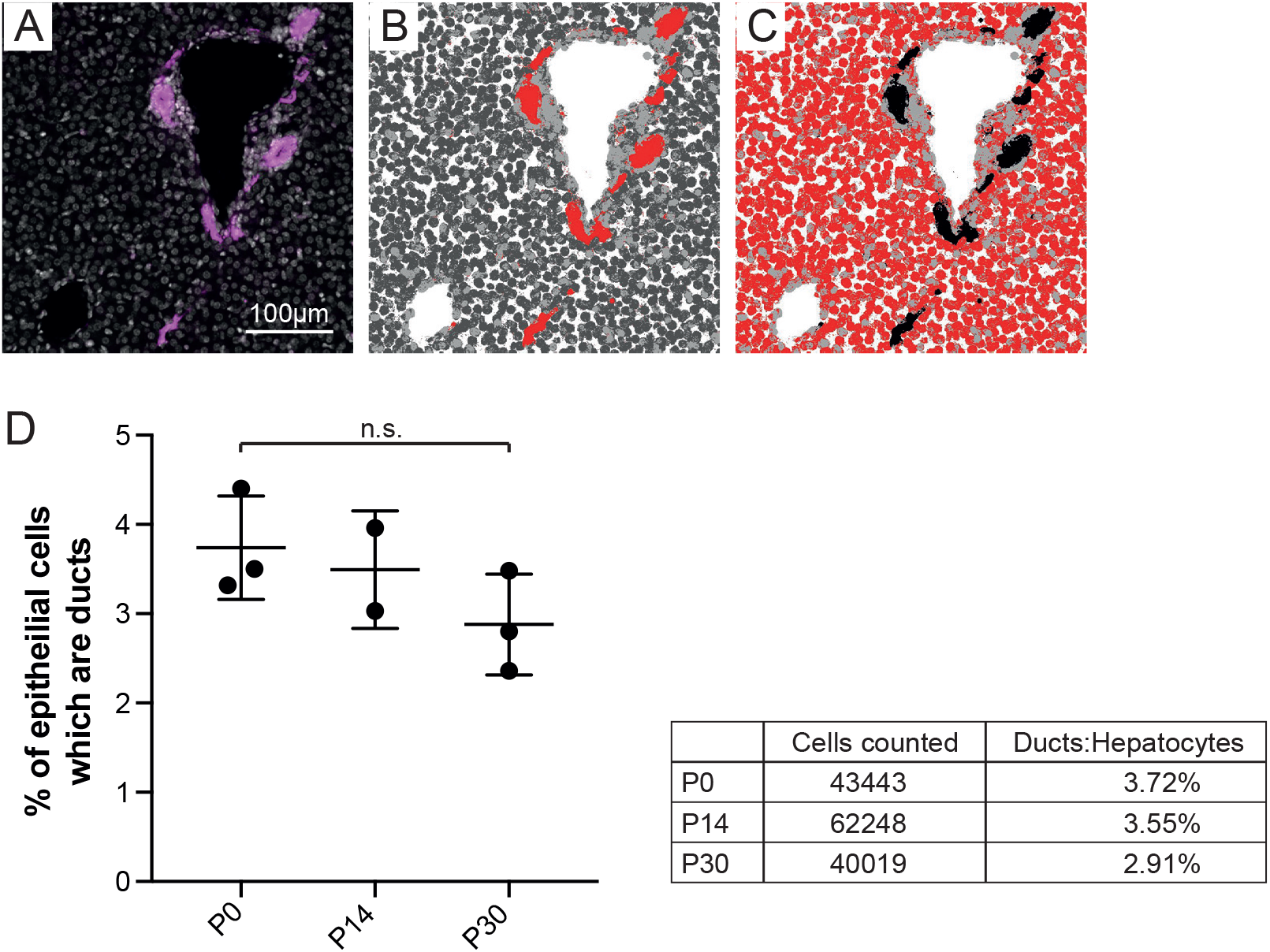
Counting of hepatocyte and ductal cell proportions in the homeostatic postnatal liver. (A-D) To count the numbers of hepatocyte and ductal cell homeostatic proportions at the different postnatal time points of interest osteopontin (OPN) staining was performed to mark the ductal cells and then the proportion of hepatocytes vs ductal cells was counted using an automated system. (A) Osteopontin (OPN, purple) marks ductal cells in the P14 liver (nuclei are counter stained with Hoecsht 33342). (B-C) Immunofluorescent images were segmented using ilastik-1.2.2 software in order to isolate cholangiocytes (B) and hepatocytes (C) (marked in red). (D) The segmented images were then imported into Fiji in order to count the number of cells using the find maxima function on the Hoechst channel. Automated counting reveals the homeostatic number of ductal cells as a percentage of epithelial cells is ~3% at all 3 time points (P0, P14 and P30) analysed. Graph represents the percentage of ductal cells within the total epithelial cells counted at the 3 time points analysed (mean ± STDEV). On average a 3.4% ± 0.6% of epithelial cells of the mouse liver are ductal, this proportion does not change over postnatal days P0 - P30 (mean ± STDEV). Table indicates the total number of cells counted and the % of ductal cells within these at each of the 3 time points analysed.

**Supplementary Figure 7:**
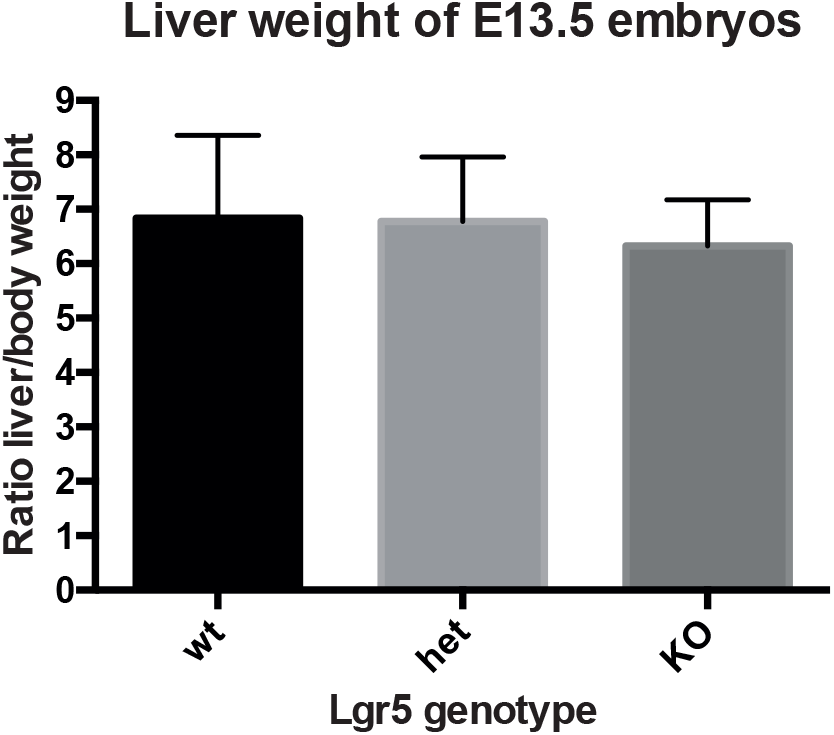
Lgr5_KO does not exhibit an apparent liver phenotype. Quantification of the ratio liver/body weight of the different genotypes for the Lgr5EGFP mouse model. Note that breeding this mouse as homozygous results in severe lethality at E17.5. Ratio liver/body weight of Lgr5EGFP WT, Het and KO mice (n=31, 5 different litters) at E13.5. No significant differences were found following analysis of all embryos or following sub-classification of genotypes according to sex (data not shown).

**Supplementary video S1: Confocal Z-stack video of a bi-potent Lgr5 clone derived from an *Lgr5-ires-creERT2;R26R-confetti* mouse embryo injected at E9.5**

**Supplementary dataset 1: scRNAseq gene lists**

**Supplementary dataset 2: Tracing counts used for the analysis**

**Supplementary dataset 3: List of material and reagents used in the manuscript**

